# Rhythmic expression of *Neurofibromin 1* in mushroom body neurons mediates circadian wake drive through activating cAMP−PKA signaling

**DOI:** 10.1101/2020.09.16.299859

**Authors:** Pedro Machado Almeida, Blanca Lago Solis, Alexis Feidler, Emi Nagoshi

**Affiliations:** Department of Genetics and Evolution, Sciences III, University of Geneva, 30 Quai Ernest-Ansermet, Geneva-4, CH-1211, Switzerland; Department of Biology, Washington and Lee University, Lexington, VA, United States

## Abstract

Various behavioral and cognitive states exhibit circadian variations in animals across phyla including *Drosophila*, in which only ∼0.1% of the entire brain neurons contain circadian clocks. This suggests that clock neurons communicate with a plethora of non-clock neurons to transmit the timing information to gate various behavioral outputs in *Drosophila*. Here, we address the molecular underpinning of this phenomenon by performing circadian RNA-seq analysis of non-clock neurons that constitute the mushroom body (MB), the center of information processing and sleep regulation. We identify hundreds of genes rhythmically expressed in the MB, including the *Drosophila* ortholog of *Neurofibromin 1* (*Nf1*), the gene responsible for neurofibromatosis type 1 (NF1). Rhythmic expression of *Nf1* promotes daytime wakefulness by activating cAMP−PKA signaling and increasing excitability of the MB. These findings reveal the pervasive, non-cell-autonomous circadian regulation of gene expression in the brain and its role in sleep, with implications in the pathology of NF1.

## INTRODUCTION

Numerous cognitive, sensory, behavioral, and emotional states exhibit circadian (near 24 h) rhythms in most animal species (Chouhan et al., 2015; Fisk et al., 2018; Lyons and Roman, 2009; McClung, 2013; Smarr et al., 2014). In humans, misalignment between day−night cycles and internal circadian rhythms due to modern lifestyle choices, such as shift work and social jetlag, correlates with a greater risk of disturbances in sleep, cognitive functions and mental health (Benca et al., 2009; Wulff et al., 2010). Although the basic mechanisms of circadian clocks and neurological underpinning of various brain functions have been extensively studied, how the former regulates the latter remains incompletely understood.

*Drosophila melanogaster* offers an unusual opportunity to study circadian regulation of brain function. The fly brain, consisting of approximately 150,000 neurons and 15,000 glia (Kremer et al., 2017), is numerically simpler and tractable with a battery of genetic tools, but shares many features with the mammalian central nervous system (CNS) (Venken et al., 2011). Flies and mammals also share the design principle and some of the components of the molecular clocks, which consist of interlocked transcriptional feedback loops. In the main feedback loop of the *Drosophila* clock, the heterodimeric transcriptional activator CLOCK/CYCLE (CLK/CYC) drives the expression of the *period* (*per*) and *timeless* (*tim*) genes. The PER and TIM proteins accumulate and dimerize in the cytoplasm, subsequently translocating into the nucleus and inhibiting CLK/CYC activity. In addition to this negative feedback loop, PDP-1 and VRILLE (VRI) regulate the transcription of *Clk*, thereby forming a stabilizing loop (Hardin, 2011). Furthermore, many features of the network organization and function of the mammalian central pacemaker, the suprachiasmatic nucleus (SCN), show similarities with the fly pacemaker neurons, such as the importance of peptidergic signaling (Renn et al., 1999) and cAMP (Levine et al., 1994). These similarities in molecular and neuronal mechanisms suggest that studies in flies can shed light on the mechanistic principles of the circadian system in mammals (Welsh et al., 2010).

Flies display circadian rhythms in a variety of behavior and brain function, such as locomotor activity, learning and memory, sleep−wake cycles, feeding, and courtship and mating (Allada and Chung, 2010); different brain areas carry out these functions. Importantly, unlike mammals where virtually all cells have clocks and can cell- and tissue-autonomously generate rhythms, within the fly CNS only approximately 150 circadian pacemaker neurons and 1,800 glia harbor molecular clocks (Hall, 2003). Therefore, circadian pacemaker neurons not only control rhythmic locomotor behavior but presumably input time-of-day information to non-clock cells in different brain areas to modulate their function and output in a circadian fashion. This regulation could occur via direct neuronal connections or indirectly through interneurons or hormonal/peptidergic signaling.

Recent studies have identified several non-clock neurons that relay the activity of pacemaker neurons to the circuits controlling locomotor rhythms and sleep−wake cycles. One locomotor output pathway involves neurons expressing the neuropeptide, leucokinin (LK) (Cavey et al., 2016). Another route includes neurons expressing diuretic hormone 44 (DH44) located in the pars intercerebralis (PI) (Cavanaugh et al., 2014). Additionally, a cluster of dopaminergic neurons, PPM3-EB, was shown to relay the activity of the pacemaker neurons to the premotor center, the ring neurons of the ellipsoid body (EB) (Liang et al., 2019). The ring neurons of the EB also play an essential role in sleep homeostasis (Donlea et al., 2018), receiving input indirectly from a subgroup of pacemaker neurons, dorsal neuron 1 (DN1), via a class of tubercular−bulbar (TuBu) neurons (Guo et al., 2018; Lamaze et al., 2018). Another subclass of pacemaker neurons, lateral posterior neurons (LPNs), is presynaptic to the dorsal fan-shaped body (dFB) of the central complex. LPNs express the allatostatin-A (AstA) neuropeptide as well as glutamate, activating dFB neurons to promote sleep (Ni et al., 2019).

There are potentially many more brain regions that are under the control of the circadian pacemaker circuit, including the mushroom body (MB).

The MB is a symmetric structure composed of ∼2,000 intrinsic neurons called Kenyon cells (KCs),the dendrites of which form a calyx, and their axonal projections form a bundle called the peduncle. At the end of the peduncle, KC axons bifurcate to form the vertical (α and α’) and medial (β, β’, and γ) lobes. Three major subtypes of KCs form these lobes: one that projects to the γ lobe and two that have branched axons projecting into the α/β lobes or α’/β’ lobes. The MB lobes can be further divided into anatomically and functionally different layers and regions (Aso et al., 2014; Keene and Waddell, 2007; Strausfeld et al., 2003). The MB is a major center of information processing in insects, analogous to the cerebral cortex, hippocampus, and piriform cortex in the human brain (Stopfer, 2014). Overwhelming evidence establishes the MB as a center for associative learning and memory (Zars et al., 2000) and regulation of sleep (Guo et al., 2011; Joiner et al., 2006; Pitman et al., 2006). Both of these behavioral processes display daily rhythms (Dubowy and Sehgal, 2017; Page, 2015). A previous study found that the main pacemaker neurons of the circadian circuit, the small ventral lateral neurons (s-LNvs), make physical contacts with some of the KCs (Gorostiza et al., 2014); therefore, it is assumed that input from the pacemaker neurons rhythmically modulates MB physiology and their functional output. However, the underlying mechanisms are unknown.

To broadly explore the mechanisms underlying the circadian modulation of MB function, we hypothesized that (1) many, if not all, MB neurons exhibit circadian rhythms in gene expression, despite containing no molecular clocks, and (2) some of these oscillating genes mediate rhythmic function of the MB. The present study tests these hypotheses by conducting circadian RNA-seq analysis of MB neurons, followed by functional analyses of candidate genes with respect to sleep, a process regulated by the MB and occurring with circadian periodicity. We identify, for the first time, a large number of genes expressed with ∼24-h period in the MB both under light-dark (LD) cycles and in constant darkness (DD). *Neurofibromin 1* (*Nf1*), the *Drosophila* ortholog of a tumor suppressor linked to neurofibromatosis type 1 (NF1), and *Pka-C1* encoding the catalytic subunit of cAMP-dependent protein kinase, are among the rhythmically expressed genes in the MB. We further demonstrate that NF1 promotes daytime wakefulness via the activation of cAMP−PKA signaling, increasing the excitability of MB neurons. Rhythmic expression of *Nf1* and *Pka-C1* gates the higher propensity for wakefulness during the daytime. Sleep is thought to be regulated by the interaction between two processes: the homeostatic sleep drive (Process S) and the circadian process driving wakefulness (Process C) (Borbely, 1982). These findings reveal the pervasive, non-cell-autonomous circadian regulation of gene expression in the brain and exemplify its role in the Process C of sleep regulation.

## RESULTS

### Circadian gene expression profiling of MB neurons

We performed circadian RNA-seq analysis of MB neurons labelled with GFP using a pan-MB neuron driver, *OK107-GAL4* (Aso et al., 2009). Brains were dissected and dissociated every 4 h from flies kept under a 12-h light −dark (LD) cycle and on the second day of constant darkness (DD) following LD-entrainment. On average, 27,000 GFP-positive cells were FACS-sorted at each timepoint, from which poly-A-tailed RNAs were amplified and sequenced following an established protocol (Abruzzi et al., 2015) (Figure 1A, Supplemental Data 1). Analysis of the RNA-seq data from biological duplicates using the JTK_CYCLE algorithm (Hughes et al., 2010) identified hundreds of genes expressed rhythmically with a ∼24-h period (Figure 1B and C). A total of 832 genes during LD and 1144 genes during DD were found to be rhythmically expressed, among which 117 genes cycled in both LD and DD (Figure 1D). These are the most conservative estimates, since analysis using the less stringent RAIN (Thaben and Westermark, 2014) and ORIOS (Larriba et al., 2016) algorithms found ∼1200 and ∼2000 genes, respectively, cycling in both LD and DD (data not shown).

**Figure 1.**
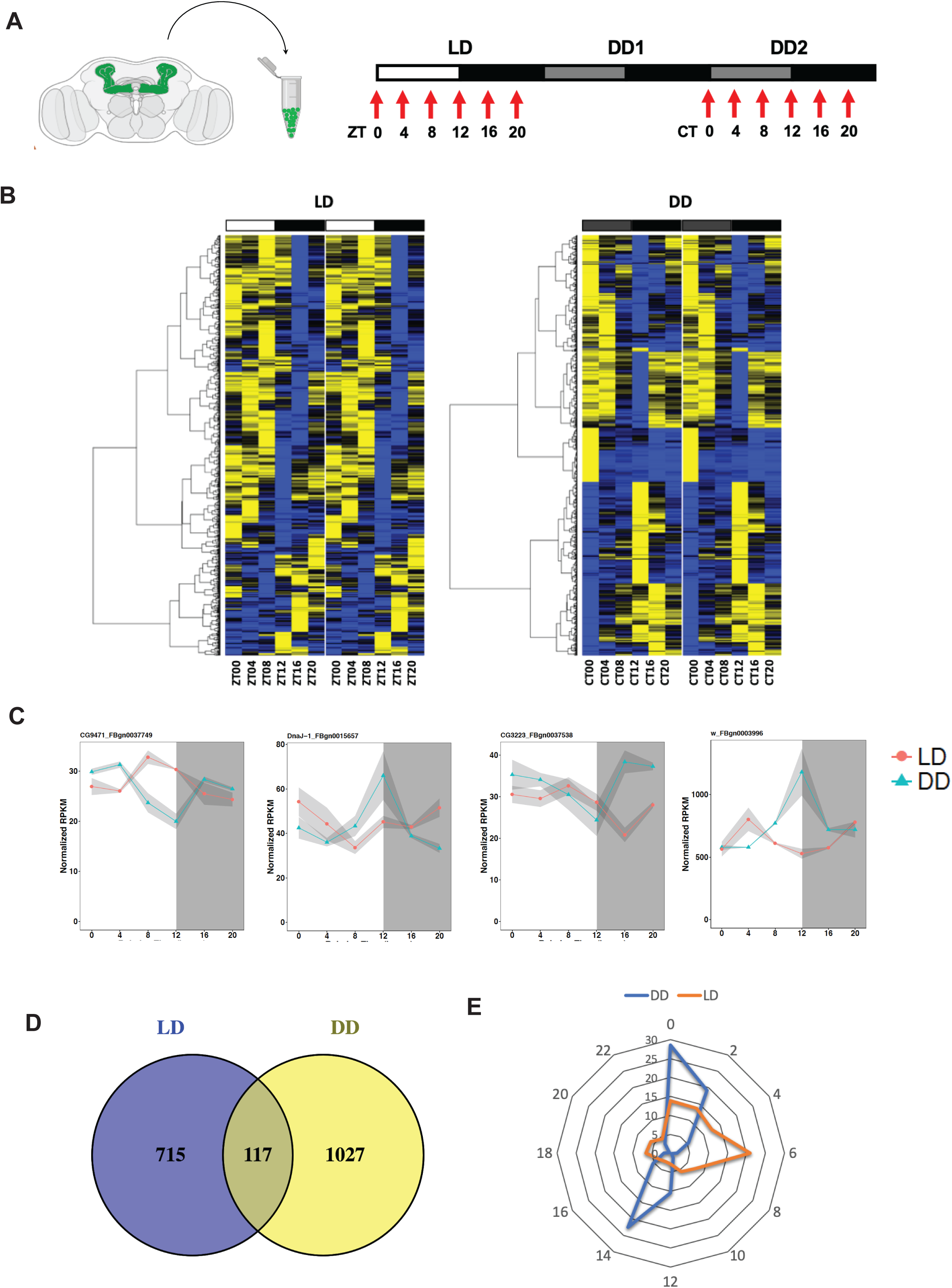
Circadian gene expression profiling of MB neurons. **A**. Schematic representation of the circadian MB neuron cell-sorting RNA-seq assay. **B**. Heatmap depicting the genes expressed with circadian oscillations during LD and DD in the MB as identified using JTK_cycle with a cut-off of *p*.*adj* < 0.05. Each row represents a gene, and rows are ordered by hierarchical clustering using the ward.D2 method. Yellow and blue colors indicate high and low relative expression levels, respectively. **C**. Examples of RNA-seq profiles of cycling genes in LD (red) and DD (blue). X-axis indicates ZT or CT. The values represent normalized RPKM. Upper and lower brackets represent the two values from the duplicates, and the dot represents the mean value. **D**. Venn diagram depicting the number of genes expressed rhythmically during LD and DD in the MB. **D**. Circular plot representing the peak phases of circadian gene expression in LD (red) and DD (blue). The number on each circle indicates the % of cycling genes peaking at a given ZT or CT.

Importantly, our RNA-seq data show the absence of canonical clock gene expression in MB neurons (Supplemental Data 1), which is evidence that rhythmic gene expression in the MB in DD is non-cell-autonomously controlled by the circadian clocks located elsewhere. The distribution of peak phases was markedly different between the LD and DD conditions: the majority of genes expressed rhythmically in LD (LD cyclers) peaked in the daytime, whereas the phases of genes cycles during DD (DD cyclers) showed a bimodal distribution, peaking around subjective dawn (CT0) or dusk (CT14) (Figure 1B and E). The data also show that ∼85% of LD cyclers became arrhythmic in DD and ∼10% of DD cyclers lost oscillations in LD (Figure 1D), indicating that light–dark cycles can drive gene expression rhythms in the MB and even impede clock-driven mRNA rhythms.

Genes expressed rhythmically in both LD and DD in the MB (dubbed MB LD-DD cyclers) are clock-driven and their rhythms are not masked by LD cycles, hence may represent functionally important genes in the natural environment. We therefore chose to focus on this subset for subsequent studies, starting with a Gene Ontology (GO) term enrichment analysis (Table S1). Despite this subset containing only 117 genes, the analysis found enrichment of several GO terms related to the processes known to be regulated by the MB, such as cognition, learning, and memory. Incidentally, these processes undergo circadian modulations (Chouhan et al., 2015; Fropf et al., 2014; Lyons and Roman, 2009). GO terms related to aminergic neurotransmitter loading and amine transport were also found to be enriched, implicating aminergic transmission in circadian rhythms of MB functions. GO terms in RNA metabolism and transcription initiation were also over-represented, suggesting the transcriptional and RNA-mediated regulation of circadian MB functioning.

### Identification of enriched *cis*-regulatory elements in the MB LD-DD cyclers

Rhythms of mRNA levels can be controlled by a transcriptional or a post-transcriptional mechanism, or a combination of both. To explore whether either mechanism is involved in cycling gene expression in MB neurons, we first searched for transcription factor binding sites (TFBSs) that are over-represented in the promoter regions of the MB LD-DD cyclers as compared with any other *Drosophila* gene using the oPOSSUM-3 tool (Kwon et al., 2012). A total of 117 TFBSs were found to be enriched in the input dataset, among which 8 exceeded the Z-score threshold (> 5.5), 11 exceeded the Fisher score threshold (> 0.55), and only 4 exceeded both; the binding sites of Optix, Twist, Onecut, and Dorsal (Table 1). This relatively restricted number of enriched TFBS motifs suggests that many cycling genes could be transcriptionally co-regulated by the same transcription factor.

**Table 1.** Enriched transcription factor binding sites in the promoters of the MB LD-DD cyclers. Top 30 transcription factor (TF) binding sites over-represented in the promoter regions of the **MB LD-DD** cycling genes, sorted by the Z-score. Significantly enriched TF binding sites that **fulfil** both a Z-score > 5.5 and a Fisher score > 0.5 are marked in bold.

| TF | JASPAR ID | Class | Family | Z-score | Fisher score |
| --- | --- | --- | --- | --- | --- |
| **Optix** | MA0199.1 | Helix-Turn-Helix | Homeo domain | **7.245** | **1.414** |
| **twi** | MA0249.1 | Zipper-type | Helix-Loop-Helix | **6.817** | **0.646** |
| **onecut** | MA0235.1 | Helix-Turn-Helix | Homeo | **6.159** | **0.577** |
| hb | MA0049.1 | Zinc-coordinating | β-β-α-zinc finger | **5.922** | 0.132 |
| **dl_1** | MA0022.1 | Ig-fold | Rel | **5.91** | **1.493** |
| br_Z1 | MA0010.1 | Zinc-coordinating | β-β-α-zinc fingerfinger | **5.853** | 0.217 |
| fkh | MA0446.1 | Winged Helix-Turn-Helix | Forkhead | **5.667** | 0.349 |
| z | MA0255.1 | Helix-Turn-Helix | Zeste | **5.516** | 0.081 |
| so | MA0246.1 | Helix-Turn-Helix | Homeo domain | 4.759 | **0.599** |
| brk | MA0213.1 | Helix-Turn-Helix | Brinker | 4.534 | **1.119** |
| Six4 | MA0204.1 | Helix-Turn-Helix | Homeo domain | 3.302 | 0.328 |
| bap | MA0211.1 | Helix-Turn-Helix | Homeo domain | 3.242 | **0.769** |
| br_Z3 | MA0012.1 | Zinc-coordinating | β-β-α-zinc fingerfinger | 2.916 | 0.359 |
| Lag1 | MA0193.1 | Helix-Turn-Helix | Homeo domain | 2.849 | 0.057 |
| hkb | MA0450.1 | Zinc-coordinating | β-β-α-zinc fingerf | 2.452 | 0.196 |
| CG42234 | MA0174.1 | Helix-Turn-Helix | Homeo domain | 2.382 | 0.271 |
| usp | MA0016.1 | Zinc-coordinating | Hormone-nuclear Receptor | 2.001 | **0.565** |
| br_Z4 | MA0013.1 | Zinc-coordinating | β-β-α-zinc finger | 1.97 | 0.016 |
| Cf2_II | MA0015.1 | Zinc-coordinating | β-β-α-zinc finger | 1.846 | 0.104 |
| D | MA0445.1 | Other Alpha-Helix | High Mobility Group | 1.775 | 0.088 |
| Su(H) | MA0085.1 | Other | LAG1 | 1.705 | **0.663** |
| CG34031 | MA0444.1 | Helix-Turn-Helix | Homeo domain | 1.704 | 0.046 |
| vvl | MA0254.1 | Helix-Turn-Helix | Homeo domain | 1.623 | **0.854** |
| slp1 | MA0458.1 | Winged Helix-Turn-Helix | Forkhead | 1.571 | 0.133 |
| Hmx | MA0192.1 | Helix-Turn-Helix | Homeo domain | 1.483 | 0.063 |
| Kr | MA0452.1 | Zinc-coordinating | β-β-α-zinc finger | 1.426 | **0.715** |
| ct | MA0218.1 | Helix-Turn-Helix | Homeo domain | 1.407 | 0.018 |
| Gsc | MA0190.1 | Helix-Turn-Helix | Homeo domain | 1.357 | 0.171 |
| achi | MA0207.1 | Helix-Turn-Helix | Homeo domain | 1.297 | 0.176 |
| ovo | MA0126.1 | Zinc-coordinating | β-β-α-zinc finger | 1.291 | 0.063 |

MicroRNAs regulate the stability and translation of mRNAs and are involved in the functioning of molecular clocks and their outputs in various organisms (reviewed in (Xue and Zhang, 2018)). At least 9 miRNAs are known to regulate *Drosophila* circadian rhythms (Niu et al., 2019; Xue and Zhang, 2018), of which 6 are expressed rhythmically in the brain (Xue and Zhang, 2018). Using the microRNA.org database (Betel et al., 2010; Betel et al., 2008), we found that all 9 circadian-relevant miRNAs have putative targets within the 3’UTRs of the MB LD-DD cyclers (Table 2). 32 genes out of 117 MB LD-DD cyclers have at least one target site of a circadian-relevant miRNA. Remarkably, *Neurofibromin 1* (*Nf1*), *shaggy* (*sgg*) and *skywalker* (*sky*) possess target sites for 3 out of the 9 circadian-relevant miRNAs. These bioinformatics predictions suggest that expression or translation of some of the MB LD-DD cyclers may be regulated post-transcriptionally by miRNAs.

**Table 2.**
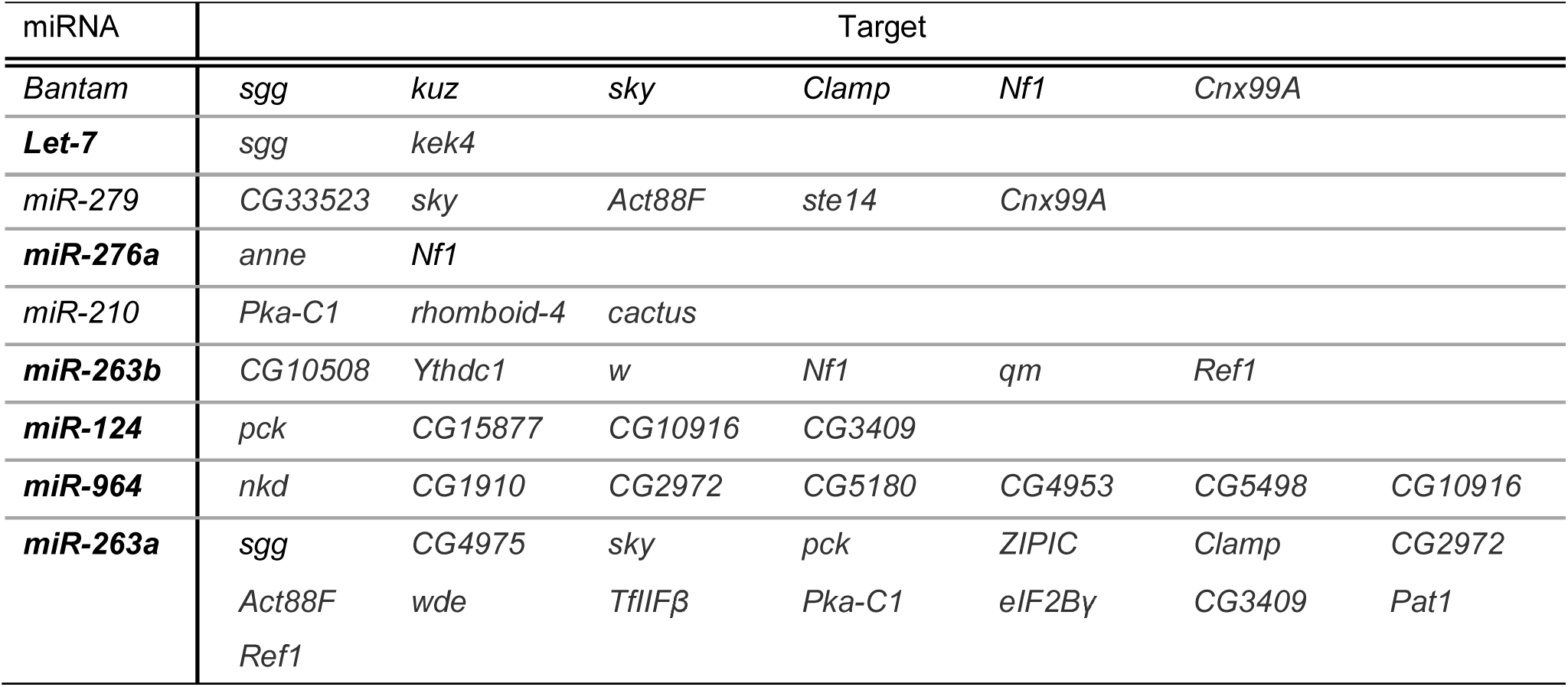
Predicted targets of circadian-relevant miRNAs among the MB LD-DD cyclers. miRNAs known to be expressed rhythmically in the fly brain are marked in bold.

### Identification of the MB LD-DD cyclers regulating sleep

Next, we asked the question whether any MB LD-DD cyclers play a role in the behavioral processes that show circadian rhythms and are regulated by the MB. We chose to focus on sleep because it is a universal behavioral state regulated by a circadian process, but the molecular aspects of circadian sleep regulation are incompletely understood. To identify genes involved in the regulation of sleep among the MB LD-DD cyclers, we generated a priority list of 21 genes based on criteria including putative functions, expression rhythm strength, and availability of fly stocks (Table S2), and knocked them down in the MB using *OK107-GAL4*. Sleep of male and female flies was measured over 5 days during LD in the *Drosophila* Activity Monitor (DAM) system and compared with controls that carry the driver only (*OK107*/+). *w*^*1118*^ was included in the assay as a reference, since most RNAi lines were generated in this background. One cross (*OK107* > *CG8735-RNAi*) did not produce viable adult flies and was excluded from this initial screen. Knockdown of 5 genes in males and 6 genes in females caused an increase in sleep (Figure 2A and S1A), but circadian locomotor rhythms remained largely normal, with the exception of *CG8142* (Table S3).

**Figure 2.**
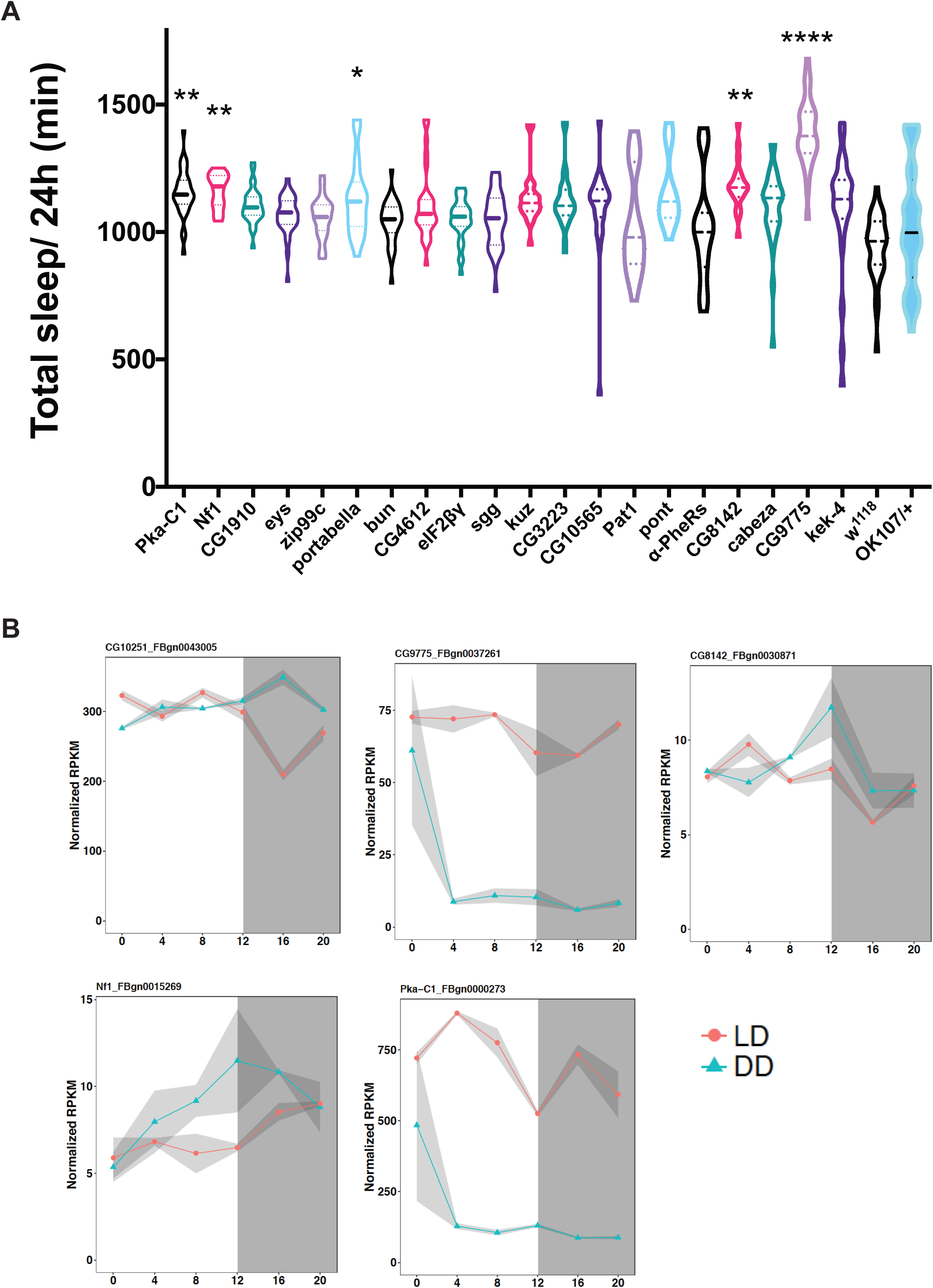
Screening for MB LD-DD cyclers involved in sleep. **A**. Violin plot of the daily total sleep in male flies expressing an RNAi transgene with the OK107 driver or *w*^*1118*^. The total amount of sleep per day was compared with *OK107/+* using the Kruskal-Wallis one-way ANOVA with Dunnett’s multiple comparisons test (\**p* < 0.05, \*\**p* < 0.01, \*\*\*\**p* < 0.0001; n = 13−32 flies per group). **B**. RNA-seq profiles of genes that increase sleep upon knockdown in the MB in males. The values are represented in normalized RPKM. Upper and lower brackets represent the two values from the duplicates, and the dot represents the mean value. RNA-seq profiles in LD and DD are displayed in red and blue, respectively.

The genes that showed a sleep phenotype upon knockdown were expressed at different levels and with different temporal patterns in the MB (Figure 2B), but all of them increased sleep. The high primary hit rate of 25% for a long-sleep phenotype is surprising and specific to the MB LD-DD cyclers, since a similar RNAi screen of 12 LD cyclers chosen according to their RNA-seq profiles identified no genes that affect sleep when knocked down in the MB (Figure S1B).

Knockdown of *Pka-C1* and *Nf1* caused an increase in sleep in both males and females (Figure 2A and S1A). *Pka-C1* encodes a catalytic subunit of cAMP-dependent protein kinase (PKA). PKA activity and sleep are inversely correlated in flies (Hendricks et al., 2001), and expression of a constitutively active form of mouse PKA-C1 (Pka-C1 mc*) (Jiang and Struhl, 1995) in the entire MB during adulthood has been shown to decrease sleep (Joiner et al., 2006). Our findings are consistent with those of previous studies and further define the role of endogenously expressed *Pka-C1* in the MB. *Nf1* is expressed in various cell types and contributes to different neuronal functions, including learning and memory (Buchanan and Davis, 2010), circadian locomotor activity (Bai et al., 2018; Bai and Sehgal, 2015; Williams et al., 2001), grooming, and sleep (King et al., 2016; van der Voet et al., 2016). Curiously, *Nf1* mutations and pan-neuronal knockdown of *Nf1* have been shown to reduce sleep (Bai and Sehgal, 2015; King et al., 2016; van der Voet et al., 2016). Our data demonstrating that *Nf1* knockdown restricted to the MB is sleep-promoting therefore suggest that NF1 can be sleep-promoting or arousal-promoting depending on the cell type.

*Nf1* encodes a large, multi-domain protein that is best characterized as a Ras-GTPase-activating protein (Ras-GAP) (Cichowski and Jacks, 2001; Yap et al., 2014). Additionally, several lines of evidence indicate that NF1 positively regulates cAMP−PKA signaling in various tissues (Hannan et al., 2006; The et al., 1997; Tong et al., 2002). *Nf1* null flies have growth defects, which can be rescued by increasing PKA activity and assimilated by inhibiting the cAMP−PKA pathway (Guo et al., 1997; The et al., 1997; Walker et al., 2006), suggesting that PKA acts downstream of, or in a parallel pathway to, NF1. Our initial results that *Pka-C1* or *Nf1* are both rhythmically expressed in the MB and wake-promoting are congruent with the convergent roles of PKA and NF1 and prompt further dissection of their functions in sleep.

### NF1 and PKA-C1 in the MB promote daytime wakefulness

We repeated the knockdown of *Nf1*and *Pka-C1* in the MB and analyzed different parameters of sleep. As in the initial screen, we drove the expression of the *Nf1* VDRC (KK) RNAi line, which has been successfully used and validated in multiple previous studies (Bai et al., 2018; King et al., 2016; van der Voet et al., 2016), with *OK107-GAL4. Nf1* knockdown profoundly increased sleep during the daytime and to a much lesser extent at night (Figure 3A and B). The increase in daytime sleep was due to a more consolidated sleep, since *Nf1*-knockdown flies had fewer sleep episodes and increased maximum sleep episode duration as compared with both driver-only and UAS-RNAi-only controls (Figure 3C−E).

**Figure 3.**
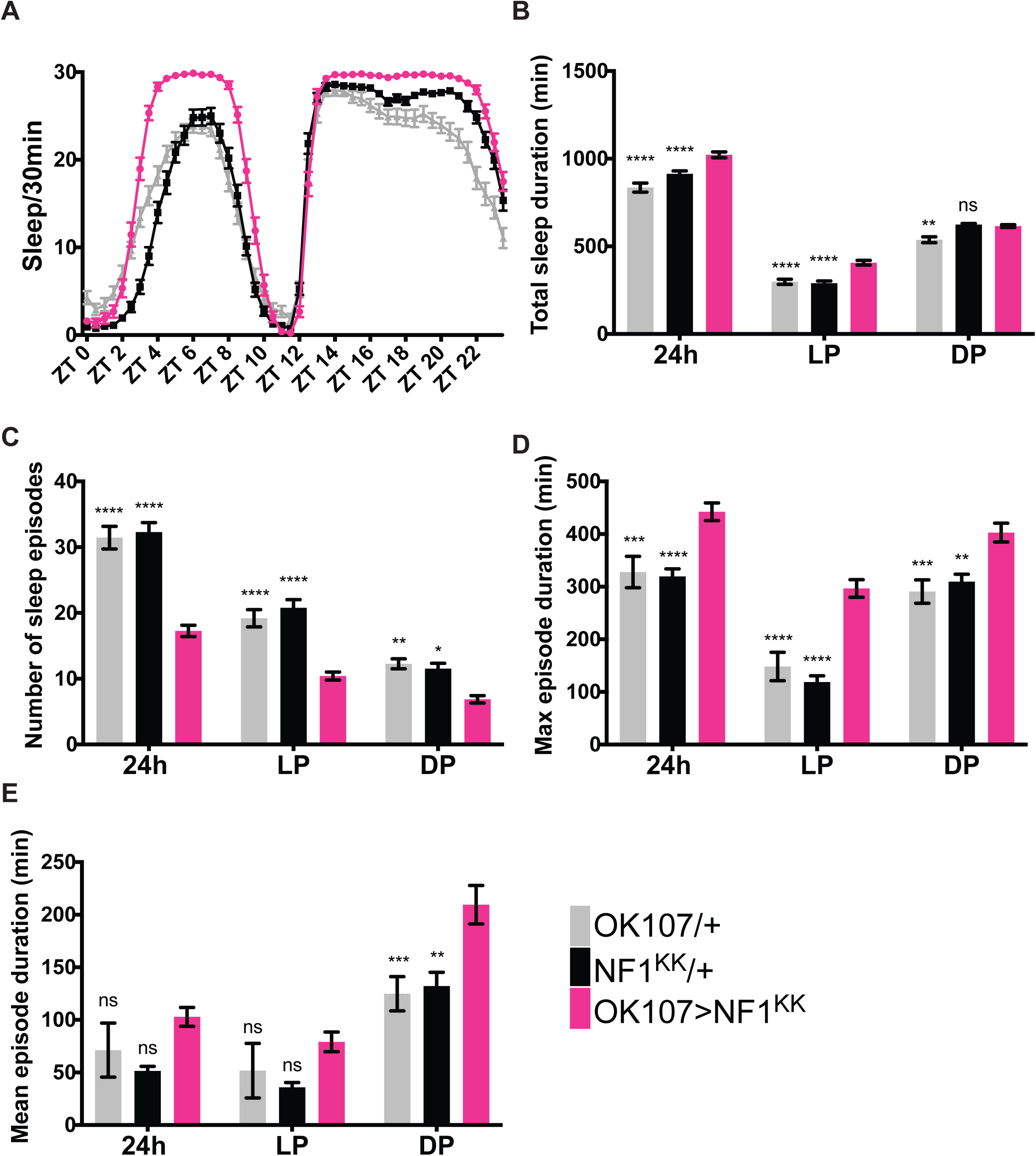
NF1 in the MB promotes daytime wakefulness. Sleep profiles of *OK107* > *Nf1*^*KK*^, GAL4-only control (*OK107*/+) and UAS-only control (*Nf1*^*KK*^/+) flies per 30-min bin averaged over 5 days during LD (n=49-59 per group). **B−E**. Different sleep variables were analyzed over a 24 h period, during the light phase (LP) and during the dark phase (DP). All values represent the mean ± SEM. Values of *OK107*> *Nf1*^*KK*^ were compared with those of *OK107*/+ and *Nf1*^*KK*^/+ using two-way ANOVA with Dunnett’s multiple comparisons test (\**p* < 0.05, \*\**p* < 0.01, \*\*\**p* < 0.001, and \*\*\*\**p* < 0.0001). **B**. Total sleep duration. **C**. Number of sleep episodes. **D**. Maximum sleep episode duration. **E**. Mean sleep episode duration.

*Nf1* mutants display growth deficiency and reduced sleep (Bai and Sehgal, 2015; King et al., 2016; The et al., 1997; Walker et al., 2006), but it remains unknown whether the growth defect is causally linked to sleep loss. To evaluate whether the sleep-promoting effect of MB-targeted *Nf1* knockdown could be developmental in origin, we restricted knockdown in the MB to adulthood using the drug-inducible MB-GeneSwitch (MB-GS) driver (Mao et al., 2004). Knockdown was induced only during adulthood by feeding flies with RU486 after eclosion. The results were similar to those obtained with a constitutive knockdown, showing longer and more consolidated sleep during the daytime (Figure S2A−F); therefore, NF1 expressed in the MB during adulthood mediates daytime wakefulness.

Reduction of PKA-C1 in the MB using shRNA consolidated and increased sleep, principally during the daytime (Figure 4A−E). Furthermore, expression of a dominant-negative mutant form of the PKA regulatory subunit PKA-R1(BDK) (Ohlmeyer and Kalderon, 1997) using OK107 also increased daytime sleep (Figure 4F−J). These results indicate that the activity of endogenous PKA-C1 in the MB promotes daytime wakefulness.

**Figure 4.**
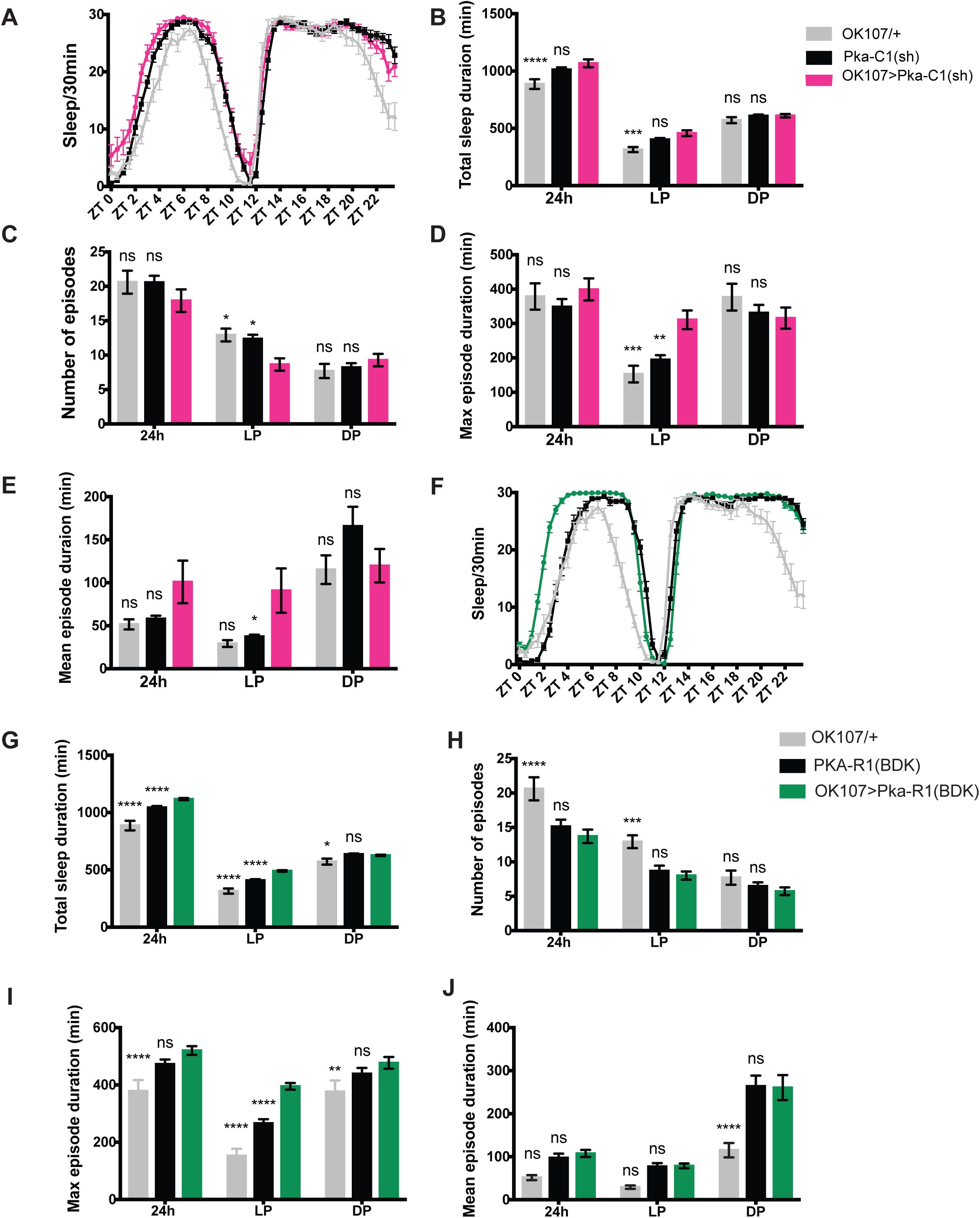
PKA-C1 activity in the MB promotes daytime wakefulness. **A−E**. Sleep profiles, amount, and structure of *OK107* > *Pka-C1*(*sh*), GAL4-only control (*OK107*/+), and UAS-only control (*Pka-C1*(*sh*)/+) flies (n=21**−**32 per group). All values represent the mean ± SEM. Values of *OK107* > *Pka-C1*(*sh*) were compared with those of *OK107*/+ and *Pka-C1*(*sh*)/+ using two-way ANOVA with Dunnett’s multiple comparisons test (\**p* < 0.05, \*\**p* < 0.01, \*\*\**p* < 0.001, and \*\*\*\**p* < 0.0001). **A**. Sleep per 30-min bin averaged over 5 days during LD. **B**. Total sleep duration. **C**. Number of sleep episodes. **D**. Maximum sleep episode duration. **E**. Mean sleep episode duration. **F−J**. Sleep profiles and analysis results of *OK107* > *PKA-R1*(*BDK*), *OK107*/+, and *PKA-R1*(*BDK*)/+ flies (n = 14−30 per genotype). Values of *OK107* > *PKA-R1*(*BDK*) were compared with those of both controls using two-way ANOVA with Dunnett’s multiple comparison’s test (\*\**p* <0.01, \*\*\**p* < 0.001, and \*\*\*\**p* < 0.0001).

Our observation that *Nf1* and *Pka-C1* are both expressed in the MB and mediate daytime wakefulness are in accordance with the idea that PKA-C1 acts downstream of, or parallel to, NF1. NF1 appears to control cAMP homeostasis through both Ras-dependent and -independent modes, although the underlying molecular mechanisms are unclear (Guo et al., 1997; Hannan et al., 2006). Thus, we sought to test whether Ras signaling mediates the effect of MB NF1 on sleep. Since expression of the constitutively active form of Ras, Ras^V12^, driven by OK107-GAL4 was lethal, we conditionally expressed it during adulthood using MB-GS (Figure S3A−E). This manipulation resulted in an overall increase in the mean sleep duration and consolidation, although to a much lesser extent than NF1 or PKA-C1 knockdown and PKA-C1 dominant negative expression, and not specific to daytime. Taken together, these results suggest that NF1 in the MB promotes wakefulness, at least partly, by a Ras-independent mechanism through the activation of cAMP−PKA signaling.

### NF1 and PKA-C1 in the MB mediate circadian wake drive

Loss of function of NF1 or PKA-C1 both increased sleep, chiefly during the daytime. We therefore wondered whether circadian expression of these genes defines a temporal pattern of their actions. To test this idea, we drove the expression of PKA-C1 mc*, a constitutively active form of the PKA catalytic subunit, with OK107. Consistent with a previous report (Joiner et al., 2006), *OK107*> *Pka-C1 mc\** led to a dramatic decrease in sleep at night by shortening sleep episodes, but did not affect daytime sleep (Figure 5B, D, F, and H). This observation suggests that endogenous PKA activity is at its highest during the daytime in the MB, and therefore constitutive activation of PKA over a period of 24 h deconsolidates nighttime sleep without further decreasing daytime sleep. The RNA-seq profile of *Pka-C1*, which peaks during the daytime, supports this interpretation (Figure 2B). Moreover, we constitutively overexpressed NF1 in the MB with OK107, which resulted in the deconsolidation of nighttime sleep, consisting of a significantly reduced mean and maximum sleep episode duration, without affecting daytime sleep (Figures 5A, C, E, and G). One can therefore infer that the activity of NF1 is also higher during the day than during the night.

**Figure 5.**
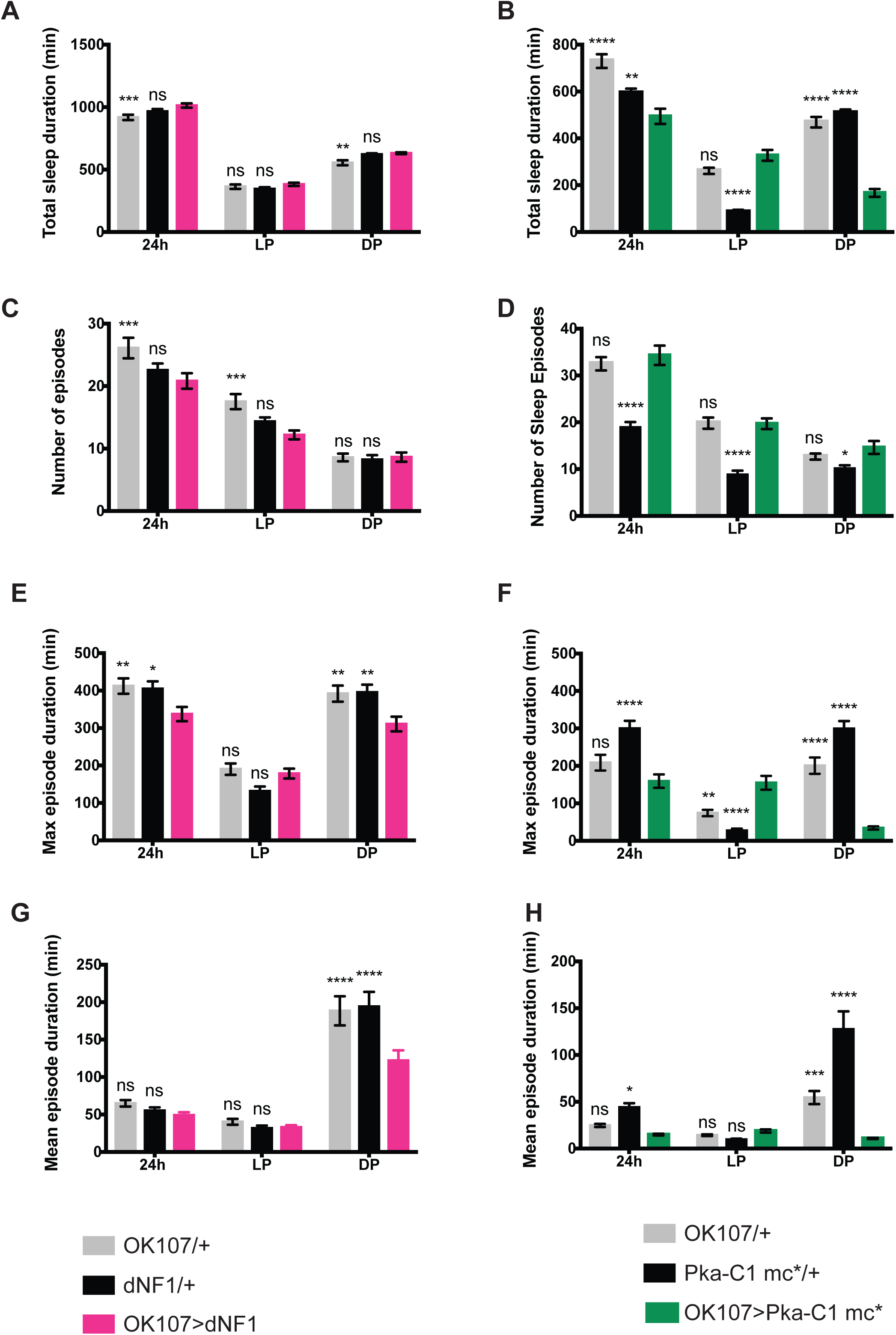
Constitutive activation of *Nf1* and cAMP−PKA signaling in the MB deconsolidates nighttime sleep. Different parameters of sleep in *OK107 > dNF1*, GAL4-only control (*OK107*/+), and UAS-only control (*dNF1*/+) (**A, C, E, and G**), *OK107* > *Pka-C1 mc\**, and its controls (*OK107*/+ and *Pka-c1 mc\**/+) (**B, D, F, and H**) (n = 50−56 per group in the *OK107* > *dNF1* experiment; n = 27−32 per group in the *OK107* > *Pka-C1 mc\** experiment). All values represent the mean ± SEM. Values of the overexpressed strains were compared with those of GAL4-only and UAS-only controls using two-way ANOVA with Dunnett’s multiple comparisons test (\**p* < 0.05, \*\**p* < 0.01, \*\*\**p* < 0.001, and \*\*\*\**p* < 0.0001). **A and B**. Total sleep duration. **C and D**. Number of sleep episodes. **E and F**. Maximum sleep episode duration. **G and H**. Mean sleep episode duration.

Since *Nf1* mRNA rhythms peak at the end of the night in LD (Figure 2B), NF1 protein levels likely peak several hours later during the daytime, driving higher wake propensity during the day than at night. Taken together with the fact that NF1 is a positive regulator of cAMP−PKA signaling, these results suggest that NF1 and PKA-C1 mediate circadian wake drive (Figure 6E).

**Figure 6.**
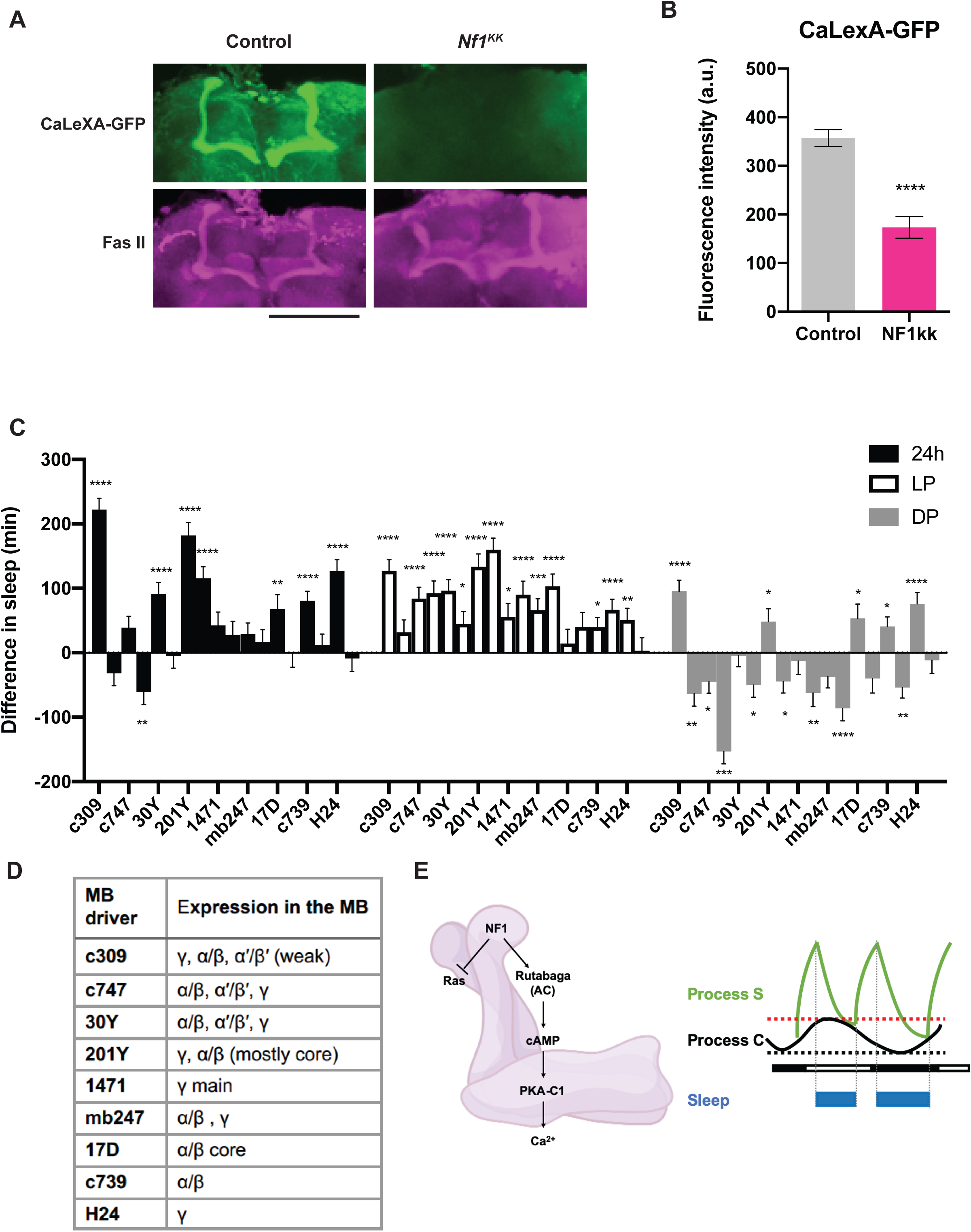
Activation of α/β and γ KCs by NF1 promotes daytime wakefulness. **A and B**. Calcium-dependent expression of GFP in the MB was monitored by driving CaLexA-GFP with *30Y-GAL4*, with or without *UAS-Nf1*^*KK*^, and co-staining with anti-GFP and anti-Fas II antibodies at ZT6. **A**. Representative confocal images. Green, CaLexA-GFP; magenta, anti-Fas II. **B**. Average GFP intensity ± SEM across all MB lobes. Values of the *30Y > Nf1*^*KK*^ were compared with those of the *30Y/+* control using the Mann-Whitney test (\*\*\*\**p* < 0.0001; n = 52 for the control, n=35 for *Nf1*^*KK*^). **C**. Effects on sleep induced by *Nf1* RNAi driven by different MB drivers. The plot represents sleep gain or loss in minutes of the experimental group (*GAL4 > Nf1*^*KK*^) as compared with the first (driver/+, left column) or second (*Nf1*^*KK*^/+, right column) control group. All values represent the mean ± SEM. Statistical analysis was performed on raw values using two-way ANOVA with Dunnett’s multiple comparisons test (\**p*<0.05, \*\**p* <0.01, \*\*\**p* < 0.001, and \*\*\*\**p* < 0.0001; n=26−87 per group). **D**. List of MB drivers and their expression patterns. **E**. A model of the role of NF1 in promoting daytime wakefulness. Left, NF1 activates PKA via Ras-dependent and - independent pathways. The Ras-independent pathway likely involves activation of adenylyl cyclase rutabaga. PKA-C1 levels and activity are higher during the daytime than at night, which gates its effect during the daytime. NF1 pathway activation increases neuronal excitability and calcium levels. Robust daytime wakefulness requires NF1 cascade activation in α/β and γ KC cells. Right, the activity of NF1 and cAMP−PKA signaling exhibit circadian rhythms that are higher during the daytime than at night. This conveys the circadian wake drive, known as Process C, in the two-process model of sleep regulation. Homeostatic sleep drive (sleep debt), Process S, increases during wakefulness and declines during sleep. Since flies display bimodal activity patterns, fly Process S is assumed to be bimodal (Geissmann et al., 2019). Process S triggers sleep near its peak and induces awakening at its trough. Circadian wake drive offsets the increase in sleep debt build-up. Knockdown of NF1 or PKA-C1 decreases circadian wake drive, thereby increasing the propensity of sleep during the daytime. Knockdown does little to nighttime sleep, since wake drive is already low (black dotted line). Constitutive expression/activation of NF1 or PKA-C1 clamps the wake drive at its maximum over 24 h (red dotted line), which increases the propensity of wakefulness during the night but has little effect on daytime sleep.

### NF1 increases the activity of MB neurons

How do NF1 and PKA-C1 in MB neurons promote daytime wakefulness? A prior study showed that cAMP−PKA activity increases intracellular calcium levels in MB neurons (Pavot et al., 2015); thus, we hypothesized that NF1 increases cAMP levels and stimulates PKA activity, thereby increasing calcium levels in MB neurons. We tested this hypothesis using the CaLexA system, in which calcium-dependent nuclear import of the LexA-VP16-NFAT fusion transcriptional activator initiates the expression of a LexAop reporter. The CaLexA system in combination with a select GAL4 driver and LexAop-GFP reports the relative calcium levels in cells targeted by GAL4 as GFP intensity (Masuyama et al., 2012). We drove CaLexA with the 30Y driver, which is expressed in all MB lobes (Aso et al., 2009) with and without *UAS-Nf1*^*KK*^. Brains of 5 to 6-day-old flies were co-immunostained for GFP and the MB marker, fasciculin II (Fas II). The CaLexA-GFP signal was significantly reduced in the presence of *Nf1* RNAi in all MB lobes, but the morphology of the MB was unaffected. Therefore, constitutive knockdown of *Nf1* in the MB reduces calcium levels without affecting development or survival (Figure 6A and B). Since calcium levels generally correlate with neuronal activity, these results indicate that NF1 increases the activity of MB neurons.

### Role of α/β and γ lobes in daytime arousal promotion by NF1

Distinct regions of the MB have been shown to be either sleep-or wake-promoting when stimulated individually (Sitaraman et al., 2015), whereas the activation of the entire or subsets of KCs results in a compound effect on sleep that does not appear to be the sum of the individual effects (Guo et al., 2011; Joiner et al., 2006; Pitman et al., 2006). Our findings that NF1 increases calcium levels across all MB lobes (Figure 6A and B) and the propensity for wakefulness (Figures 3 and 5) prompted us to examine whether the wake-promoting role of NF1 is localized to specific MB subregions. To this end, we drove the expression of *Nf1* RNAi with a set of GAL4 drivers expressed in different subsets of KCs (Aso et al., 2009) (Figure 6C and D). Sleep was affected both positively or negatively by a variety of these drivers, consistent with the notion that the MB contains both sleep- and arousal-promoting neurons. Notably, however, the increase in daytime sleep, the most conspicuous phenotype induced by NF1 knockdown in the entire MB, was recapitulated with most of the drivers.These include c739, which is expressed mainly in the α/β lobe, and 1471-GAL4, which is expressed only in the γ main lobe. The increase in daytime sleep was most pronounced with 201Y, which is expressed in the α/β and γ lobes (Figure 6C, 6D, and Table S4).

To further define KC subtypes where NF1 exhibits its wake-promoting role, we drove *Nf1* RNAi with a set of MB-split GAL4 divers that target more refined KC subtypes (Table S5). No single split-GAL4 driver was sufficient to change the total sleep duration; however, *Nf1* knockdown in α/βs KCs reduced the sleep episode number and increased the maximum sleep episode duration during the daytime. Knockdown in the γ main and γ dorsal KCs also increased the daytime maximum sleep episode duration. Thus, consolidation of daytime sleep structure may be attributed to the α/βs, γ main, and γ dorsal KCs. Taken together, these results suggest that NF1 expressed in most subtypes of KCs contributes to daytime wakefulness, but the overt effect of NF1 on sleep output emerges from simultaneous activation of α/β and γ KCs.

## DISCUSSION

Here we employed circadian RNA-seq analysis to identify a large number of genes expressed with diurnal and circadian rhythms in MB neurons, which do not harbor molecular clocks. We demonstrate that some of the genes expressed with circadian patterns in the MB regulate sleep−wake cycles, specifically by promoting wakefulness. Circadian expression of *Nf1*, the *Drosophila* ortholog of the tumor suppressor *NF1*, promotes daytime wakefulness through the activation of subsets of MB neurons. The wake-promoting role of *Nf1* is mediated by the cAMP−PKA signaling, and circadian expression of *Nf1* and Pka*-C1* gates wake drive during the daytime. These findings generally support the two-process model of sleep regulation and suggest that Process C (circadian wake drive) may be partly mediated in the MB by NF1−cAMP/PKA signaling (Figure 6E)(Borbely, 1982).

### Circadian gene expression in non-clock neurons

We show that 832 genes are expressed rhythmically in LD and 1144 genes cycle on the second day during DD in MB neurons (Figure 1B−D). These numbers, i.e., nearly 10% of the genes in the *Drosophila* genome, are surprisingly close to the percentage of cycling genes in a given clock-containing cell across metazoan (Kula-Eversole et al., 2010; Panda et al., 2002; Storch et al., 2002). These results are the first to demonstrate free-running gene expression rhythms in cells without canonical molecular clocks. Circadian RNA-seq analysis in non-clock-containing, TH-GAL4-labeled dopaminergic neurons has been reported, but, only LD timepoints were analyzed, showing approximately 100 cycling genes(Abruzzi et al., 2017). This unexpected difference in the number of cycling genes between MB neurons and dopaminergic neurons suggests that the physiology of the MB is tightly controlled by the clock neurons, highlighting its significance. Our data also inform omics studies that time-of-day differences should be considered even when targeting non-clock cells.

Only ∼15% of LD cyclers and ∼10% of DD cyclers are rhythmically expressed under both LD and DD conditions. This relatively small overlap, in addition to the marked difference in phase distribution between LD and DD, indicates a strong control of gene expression in the MB by light (Figures 1B−E). These findings are congruent with the notion that the MB is directly connected to multiple primary sensory processing centers including the optic lobe (Vogt et al., 2016; Yagi et al., 2016). The MB integrates multi-modal sensory inputs, including time of day, to coherently perceive the external environment. Our results suggest that gene expression changes within the MB driven by sensory information may be part of information processing.

Light- and clock-driven genes appear to play different biological roles in the MB. This idea is supported by the results of the mini-RNAi screen, which found ∼25% of candidates from the MB LD-DD cyclers to be involved in sleep regulation but none from the LD cyclers (Figures 2A, S1A, and S1B). The screen was by no means exhaustive, since only one RNAi line was used per gene due to its exploratory nature, and had the same limitations that are common to all genetic screens. Nonetheless, the results are broadly in agreement with our assumption that clock-driven genes that keep cycling under LD cycles may carry out instructive roles in behavioral regulation, instead of merely responding to inputs from sensory systems or the circadian circuit. The diurnal environmental cycle is perceived as rhythms in the sensory modality, which potentially modulate a wide range of cognitive and behavioral states regulated by the MB. The list of cycling genes established in the present study provides a valuable resource for the identification of genes involved in such mechanisms by employing a panel of behavioral paradigms, such as associative learning assays using odor (McGuire et al., 2004; Pitman et al., 2009), visual information (Schnaitmann et al., 2010) or time of day (Chouhan et al., 2015) as conditional stimuli.

How can gene expression rhythms be driven in the cells that do not contain circadian clocks? Enrichment of binding sites for transcription factors and circadian-relevant miRNAs in cycling genes suggests that both transcriptional and miRNA-mediated mechanisms may be involved (Tables 1 and 2). The possibility of other post-transcriptional mechanisms, such as splicing, can be further tested gene-by-gene using qPCR or by sequencing total RNA in future studies. The relatively small number of enriched TFBSs could be due to the size of the input dataset (117 genes) as well as the heterogeneity of KCs. It should be noted that, several enriched TFBSs are the targets of TFs known to be expressed and to function within the MB. *Broad* (*br*) is expressed in the adult MB (Restifo and Hauglum, 1998), specifically within the α/β lobes (Shih et al., 2019). The expression of *forkhead* (*fkh*) and *ultraspiracle* (*usp*) in the adult MB have also been shown (Boulanger and Dura, 2015; Kobayashi et al., 2006). A previous study reported the enrichment of mRNA encoding *dorsal* (*dl*) in the α/βp MB neurons (Shih et al., 2019). Overexpression of *dl* in the MB leads to fewer KCs and defasciculation of their axons (Nicolai et al., 2003). Interestingly, *cactus* mRNA, which encodes a cofactor of Dorsal, is among the dataset of genes that cycle in the MB in both LD and DD (Supplementary Data 1). This suggests a possibility that a Cactus/Dorsal complex accumulates rhythmically in the MB, thereby controlling rhythmic expression of its target genes.

Some of the TFBSs identified in the cycling gene promoters did not reach the Z-score or Fisher-score cut-offs. This could be due to the heterogeneity of KCs, and they may nevertheless regulate rhythmic gene expression in a subset of KCs. For instance, Chorion factor 2 (Cf2) has been shown to rhythmically drive the expression of mir-276a, which in turn binds to the 3’UTR of *timeless* and inhibits the translation of TIM at the proper time of day (Chen and Rosbash, 2016). A similar mechanism may take place during the course of *Nf1* and *anne* expression, since they are targets of mir-276a (Table 2).

In the absence of clock gene expression within in MB neurons, it is plausible that input from the circadian system provides temporal control of these transcriptional and/or post-transcriptional mechanisms. Which cells are afferent to the MB and how do they transmit time-of-day information to modulate gene expression in KCs? The heterogeneity of KCs poses a further challenge in answering these open questions. A previous study used GRASP (GFP Reconstitution Across Synaptic Partner) (Feinberg et al., 2008) to demonstrate the physical contact between s-LNv pacemaker neurons and KCs (Gorostiza et al., 2014). Despite the fact that it remains unanswered whether the observed GRASP signal represents functional connectivity, this finding suggests a possible monosynaptic input from circadian pacemaker neurons to KCs. The circadian system may also communicate with the MB via indirect synaptic inputs, volume transmission, and long-range signaling with neuropeptides. Whole-brain connectomics (Zheng et al., 2018) may provide important information to address these possibilities.

### Role of NF1 in sleep

We demonstrate that *Nf1* in the MB is wake-promoting. Previous studies have shown an opposite, sleep-promoting role of ubiquitously expressed *Nf1* (King et al., 2016; Williams et al., 2001). Since we used the same *Nf1*^*KK*^ RNAi strain as in their studies, the phenotypic differences are not experimental artifacts but indicate that NF1 can be both wake- and sleep-promoting depending on the cell type. The bidirectional regulation of sleep may be achieved by activating or inhibiting wake- or sleep-promoting neurons. More surprisingly, whereas we found that NF1 activates MB neurons (Figure 6A and B), the Sehgal group showed that NF1 inhibits the activity of peptidergic neurons (Bai et al., 2018) using the same functional calcium imaging method. Therefore, NF1 downstream pathways also seem to diverge depending on the cell type.

In the MB, NF1 activates cAMP−PKA signaling, which is at least partly independent of Ras and leads to an increase in calcium levels (Figures S3, 6A, and 6E.). Both circumstantial and experimental evidence from previous studies support these findings. *Nf1* deficiency reduces cAMP levels in *Drosophila* embryonic brains (Tong et al., 2002) and in murine astrocyte cultures(Dasgupta et al., 2003). Fly NF1 activates AC1 encoded by *rutabaga* (Guo et al., 1997; The et al., 1997), which is initiated by its interaction with the C-terminal domain of NF1 (Hannan et al., 2006). Moreover, it has been shown that knockdown of *rutabaga* with *OK107-GAL4* reduces the nicotine-evoked Ca^2+^-response in the MB, whereas *OK107 > Pka-C1 mc\** has the opposite effect (Pavot et al., 2015). PKA likely phosphorylates K^+^ channels (Bruggemann et al., 1993; Delgado et al., 1992), thereby rendering the cells more excitable.

A previous study showed that the expression of *UAS-Pka-C1 mc\** is sleep promoting when driven by *201Y*-*GAL4* but wake-promoting when driven by *c309*-*GAL4* (Joiner et al., 2006). *201Y* is mostly expressed in the α/β core and γ neurons, while *c309* expression covers all MB lobes, similar to *OK107*. A functional calcium imaging study reported that calcium levels increase broadly in the MB when flies are awake and decrease with sleep (Bushey et al., 2015). In contrast, thermogenetic activation of individual subsets of KCs using split-GAL4 drivers has shown that α’/β’ and γ main neurons are wake-promoting and γ dorsal neurons are sleep-promoting (Sitaraman et al., 2015). Therefore, although individual KC subtypes can be sleep- or wake-promoting when stimulated separately, simultaneous and constitutive activation of most KCs causes a net increase in wakefulness in baseline sleep. This is also in contrast to the situation when the majority of KCs are temporarily activated (Kayser et al., 2014; Pitman et al., 2006). Our findings are in accordance with this interpretation. Importantly, although α/β and γ neurons are most relevant for the action of *Nf1*, our data do not exclude the contribution of other KC subtypes (Figure 6C, Tables S4 and S5).

*Nf1* mutant flies have learning disabilities (Guo et al., 2000; Ho et al., 2007), reminiscent of one of the manifestations of NF1 (Williams et al., 2009). *Nf1*-dependent olfactory learning in flies involves the cAMP pathway and requires the expression of *Nf1* in the α/β lobes of the adult MB (Buchanan and Davis, 2010). Our study shows, for the first time, a role for *Nf1* expressed in the MB in sleep and identifies the α/β lobe as a critical site for its action (Figure 6C, Tables S4 and S5). Considering that patients with NF1 have a higher prevalence of sleep disturbances, including daytime sleepiness (Licis et al., 2013; Marana Perez et al., 2015), the fact that *Nf1* in the same subset of MB neurons is necessary for both memory formation and sleep regulation in flies is remarkable. These findings also suggest that the study of the role of *Nf1* expressed in the MB may shed light on some aspects of NF1 pathology.

## ACKNOWLEDGEMETS

We thank Simon Sprecher for *MB-GS* line, Daniel Kalderon for *UAS-PKA-C1 mc\**, James Walker for *UAS-Nf1*, and Bloomington Drosophila Stock Center and VDRC for other fly strains. We also thank Giorgio Gilestro and Ueli Schibler for their helpful comments on this study. This research was funded by grants from the Swiss National Science Foundation (31003A_169548 and 310030_189169) and the JST PRESTO program. We are grateful for the support of the EuroScholors program to AF and the Plan Strategique Sciences Vie (PSVIE) of the University of Geneva to BL. Some figures were created using BioRender.com.

## AUTHOR CONTRIBUTIONS

EN and PM and planned the experiments. PM, BL, and AF conducted the experiments. PM, BL, AF and EN analyzed the data. EN wrote the manuscript.

## Methods

### Fly strains

All *Drosophila melanogaster* strains were maintained on standard cornmeal-agar food at 18°C or 25°C in a 12-h light−dark cycle and controlled humidity. Detailed information on the strains used in the present study is provided in the table below.

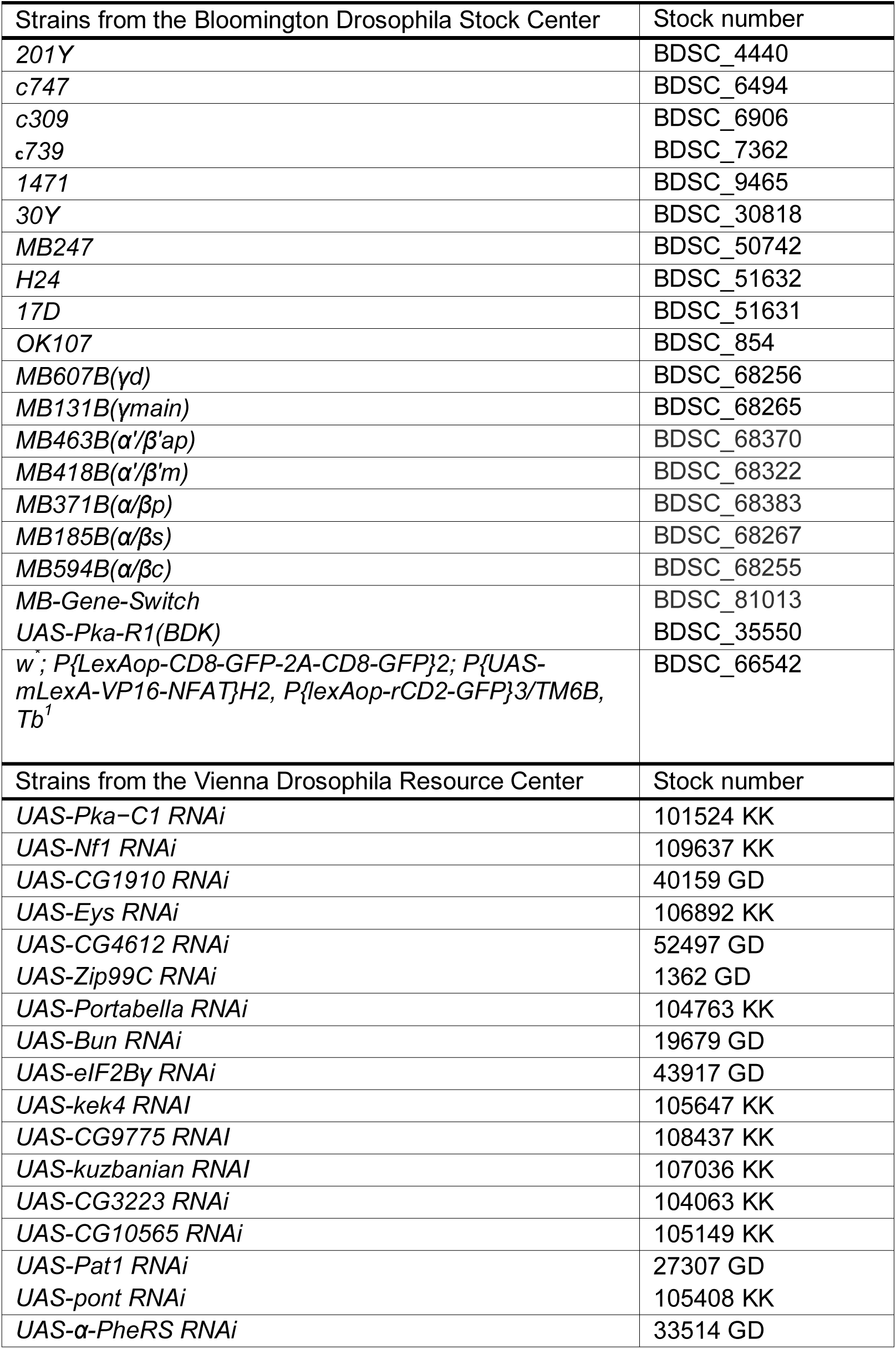
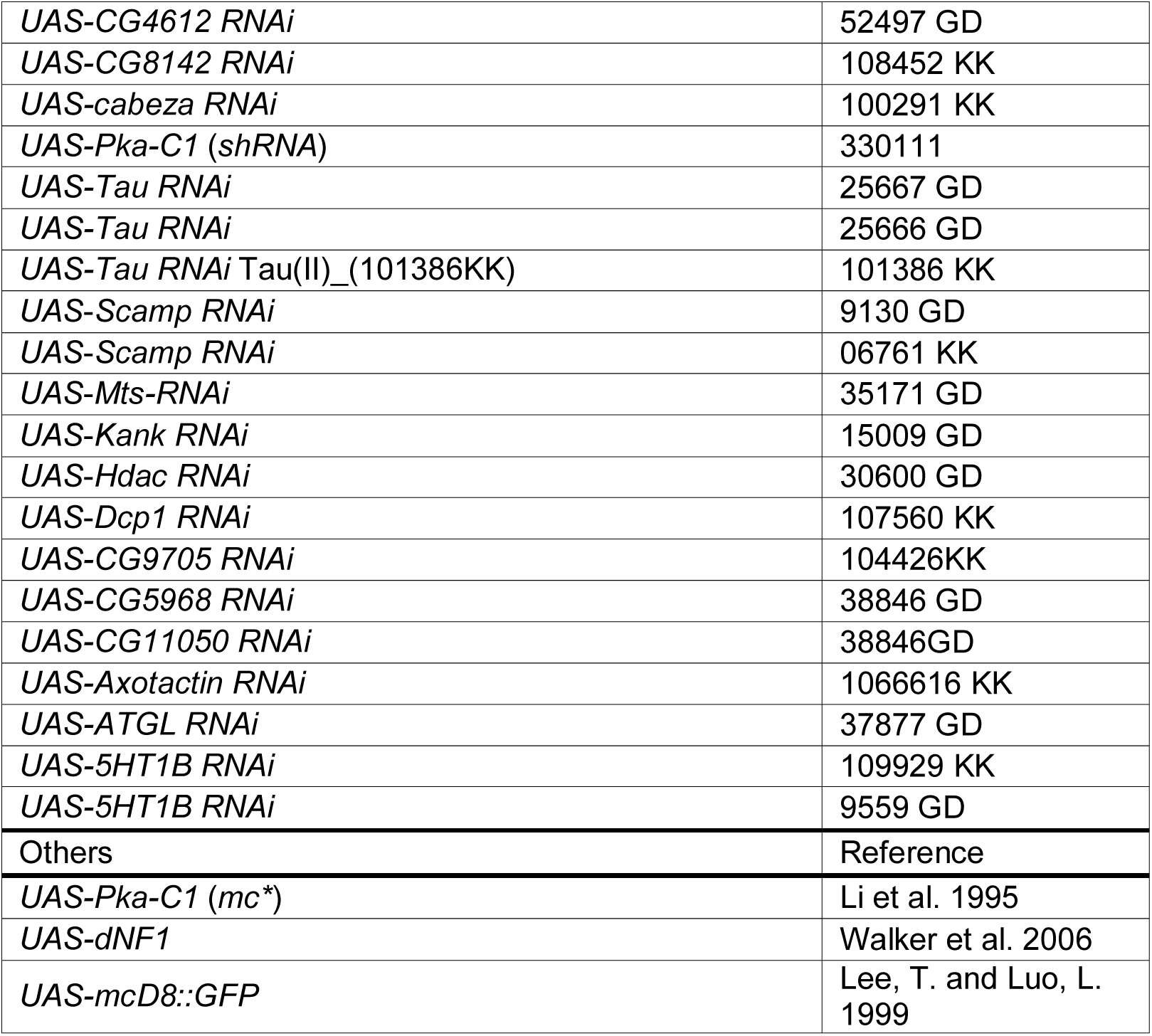

### Brain dissection and collection of MB neurons

To collect cells during the LD, 2-to 4-day-old adult flies carrying the *OK107-GAL4, UAS-mCD8::GFP* transgenes were entrained to LD cycles for 3 days and dissected on the 4th day of LD. DD samples were collected on the second day of DD following 3 days of entrainment to LD. At each timepoint, the brains of 25 flies, including both males and females, were dissected and triturated following the protocol described in (Abruzzi et al., 2015; Nagoshi et al., 2010). Optic lobes were removed and discarded during dissection. The cell suspension was kept on ice in a low-binding Eppendorf tube and immediately processed for fluorescent-activated cell sorting (FACS). Sorting of the GFP-positive cells was performed at the Flow Cytometry Core Facility of the University of Geneva using a Bio-Rad S3 Cell Sorter. All procedures were performed using low-binding plasticware. Before sorting, the cell suspension was filtered through a Filcon 70-µm strainer (BD biosciences, Cat. # 340606). Roughly 27,000 GFP-positive cells (equivalent to ∼350µL) were collected into 500µL Buffer RLT Plus (QIAGEN RNeasy Plus Micro Kit, Cat. # 74034). Midway through the sorting process, the machine was halted in order to vortex the suspension containing the first ∼13,500 collected cells. This ensures the proper lysis of the cells and the collection of cell-containing droplets that may be on the walls of the centrifuge tube. The collection tube was vortexed once more at the end of the sorting. Cells and cell lysates were kept on ice during all steps. Total RNA extraction was performed according to the manufacturer’s protocol (QIAGEN RNeasy Plus Micro Kit, Cat. # 74034) and eluted in 14µL supplied elution buffer. From the ∼12µL of total RNA collected, 1.5µL was transferred into a separate tube to test RNA integrity and concentration. All samples were flash-frozen in liquid nitrogen and stored at - 80°C until further processing.

### RNA-seq of isolated MB neurons and data analysis

All steps required for the RNA integrity check, library preparation and sequencing were performed by experts at the iGE3 Genomics Platform at the University of Geneva. Total RNA integrity and quantitation was performed using the Bioanalyzer High Sensitivity RNA 6000 Pico kit (Agilent, Cat. # 5067-1513). cDNA library preparations were conducted using the SMART-Seq v4 Ultra Low Input kit from TaKaRa (Cat. # 634894) according to the manufacturer’s instructions. Libraries were sequenced on the HiSeq2500 Illumina machines using single-end 50-bp reads to generate more than 10 million reads per library. The iGE3 Genomics Platform performed QC, trimming of sequencing adaptors, and filtering of low-quality reads. Raw reads were aligned with the Dm3 *Drosophila* genome using the EPFL’s HTSstation platform pipeline (David et al., 2014). All libraries generated on average ∼18 million single-end 50-bp reads, from which approximately 80% (∼14 million reads) were mapped to the genome. The normalized expression levels (Reads Per Kilobase of Million mapped reads; RPKM) were used for downstream analysis.

### Analysis of circadian gene expression

Genes with an expression <1 RPKM at any timepoint were excluded from the circadian analysis. Expression profiles were analyzed for rhythmicity in R (r-project.org) using JTK_CYCLE (v3.1) (Hughes et al., 2010). RNA expression profiles and heatmaps were produced using ggplot2 in R. The significance threshold was set to an adjusted *p*-value (*p*.*adj*) < 0.05.

Gene Ontology term enrichment analysis of cycling genes was performed using *GOrilla* (Eden et al., 2009). Transcription factor binding sites (TFBS) within the ± 2 kb of the transcription start site were identified using the oPOSSUM-3 tool (Kwon et al., 2012). miRNA target site analysis was performed using the microRNA.org database (Betel et al., 2010; Betel et al., 2008).

### Sleep assays and analysis

The fly sleep assay was performed using the Drosophila Activity Monitoring (DAM) system (Trikinetics) as described previously (Agosto et al., 2008). Briefly, 2-to 5-day-old male or virgin female flies were placed individually in 65 mm x 5 mm glass tubes containing 5% agarose with 2% sucrose. To induce MB-GeneSwitch expression, 200 µg/mL RU486 (Mifepristone; Sigma, Cat. # M8046) was added. Flies were entrained for 3 days in LD (12-h light−dark), and data for sleep analysis were collected during the following 5 days of LD. Locomotor activity was collected in 1-min bins, from which sleep quantity, defined as bouts of inactivity lasting for 5 min or longer (Hendricks et al., 2000; Shaw et al., 2000), was extracted. All sleep behavior assays were performed at 25°C.

Sleep analysis was performed in MATLAB (MathWorks, version R2017a, build 9.2.0.538062) using SCAMP (Vecsey laboratory; (Donelson et al., 2012)) according to the instruction manual. The raw data obtained with SCAMP were compared using ANOVA followed by Dunnett’s *post-hoc* tests for multiple comparisons in GraphPad Prism (v.8.1). Unless otherwise indicated, all sleep data were analyzed from experiments using virgin female flies and plots were generated in GraphPad Prism (v.8.1).

### Immunohistochemistry and microscopy

GFP expression in the MB driven by the CalexA system was visualized using whole-mount brain immunofluorescence with anti-GFP and anti-Fasciclin II (Fas 2) antibodies following the previously described protocol (Bou Dib et al., 2014) with minor modifications. Briefly. 5-to 6-day-old flies were decapitated at ZT6 following 3 days of LD-entrainment, and the heads were fixed in 4% paraformaldehyde + 0.3% Triton X-100 for 1 h on ice, and washed quickly twice and then twice for 20 min in PBST-0.3 (PBS, 0.3% Triton X-100). Subsequently, the head cuticle was partly removed, and the heads were washed twice more, blocked in blocking solution (55% normal goat serum, PBST-0.3) for 1 h at room temperature, and incubated with the primary antibodies for 48 h at 4°C. Rabbit affinity purified anti-GFP (1:500, Invitrogen G10362) and 1D4 mouse anti-Fasciclin II (1:5; Developmental Studies Hybridoma Bank (DSHB)) (were used as primary antibodies, and Alexa 488 goat anti-rabbit IgG (1:250, Thermo Fisher A11008) and Alexa 633 goat anti-mouse IgG (1:250, Thermo Fisher A21052) were used as secondary antibodies. The remaining cuticle was removed from the head and the brains were mounted on slides with Vectashield (Vector Laboratories). Images were acquired using a Leica TCS SP5 confocal microscope.

### Image analysis

Confocal Z-stacks were analyzed using the Image J/Fiji software (Schindelin et al., 2012). GFP fluorescence intensities in the MB were measured from the Z-projections within the area defined by the Fas II signal.

### Statistical analysis

Statistical analysis of sleep data was performed using one- or two-way ANOVA with Dunnett’s *post-hoc* multiple comparisons test in GraphPad Prism (v.8.1). Statistical comparison of fluorescence intensities was performed using the Mann−Whitney test. Significant values in all figures: \**p* < 0.05, \*\**p* < 0.01. \*\*\**p* < 0.001, and \*\*\*\**p* < 0.0001. ns indicates not significant.

## Supplemental Information titles and legends

**Supplemental Data 1**

Raw read counts and RPKM of RNA-seq data of MB neurons isolated during LD and DD, and the results of cycling expression analysis using JTK_CYCLE.

