## Supplemental files for "Rhythmic expression of *Neurofibromin 1* in mushroom body neurons mediates circadian wake drive through activating cAMP−PKA signaling"

**SUPPLEMENTAL INFORMATION**

**Supplemental Figures**

**Figure S1**

**Figure S2**

**Figure S3**

**Supplemental Table titles and legends**

**Table S1**

**Table S2**

**Table S3**

**Table S4**

**Table S5**

A

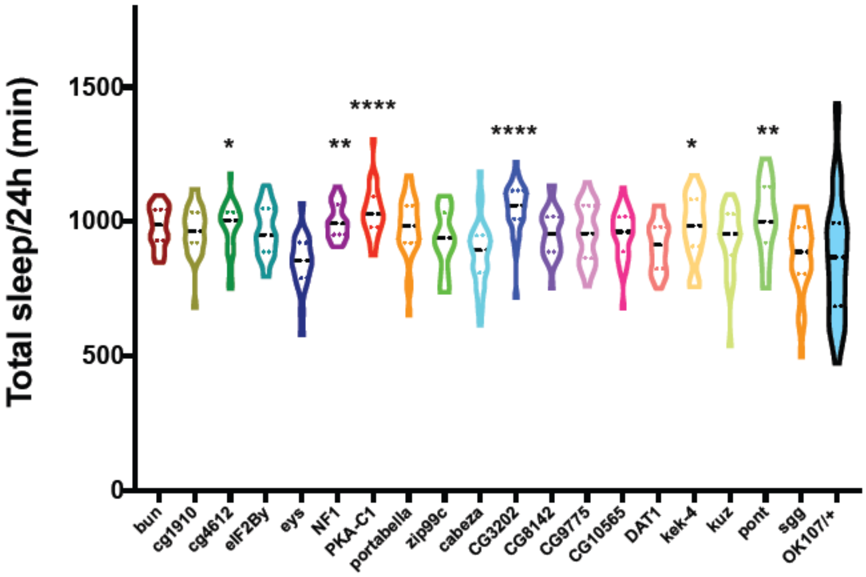


B

| Genotype | n | Total sleep/24h (min) | SEM |
|---|---|---|---|
| Tau(III)_(25667GD) | 32 | 1118.8 | 15.5 |
| Tau(II)_(25666GD) | 32 | 1088 | 17.0 |
| Tau(II)_(101386KK) | 32 | 1181.8 | 11.6 |
| Scamp(II)_(9130GD) | 20 | 1086.5 | 16.6 |
| Scamp(II)_(106761KK) | 32 | 1157.8 | 15.9 |
| Mts(III)_(35171GD) | 32 | 1095.9 | 14.1 |
| Kank(II)_(15009GD) | 31 | 1084.2 | 16.9 |
| Hdac1(II)_(30600GD) | 32 | 1080.3 | 20.1 |
| Dcp1(II)_(107560KK) | 31 | 1113.2 | 14.9 |
| CG9705(II)_(104426KK) | 31 | 1096.7 | 21.0 |
| CG5968(II)_(38846GD) | 27 | 1067.8 | 18.1 |
| CG11050(II)_(107837KK) | 32 | 1101.1 | 14.1 |
| Axotactin(II)_(1066616KK) | 22 | 1111.8 | 17.4 |
| ATGL(II)_(37877GD) | 31 | 1074.4 | 16.2 |
| 5HT1B_(109929KK) | 31 | 1163.1 | 15.1 |
| 5HT1B(III)_(9559GD) | 32 | 1141.5 | 12.6 |
| OK107/+ | 31 | 1136.1 | 13.1 |

**Figure S1. RNAi screening of the MB LD cyclers for sleep**
**A.** Violin plot of the daily total sleep of female flies expressing an RNAi transgene with the OK107 driver. The total amount of sleep per day was compared with *OK107/+* by The Kruskal-Wallis one-way ANOVA with Dunnett's multiple comparisons test (*$p < 0.05$, **$p < 0.01$, ****$p < 0.0001$; n = 21–46 flies per group). **B.** Results of the screening of the MB LD cyclers for sleep. Each gene was knocked down with the OK107 driver. No significant change in total sleep amount was detected by ANOVA with Dunnett's multiple comparisons test.

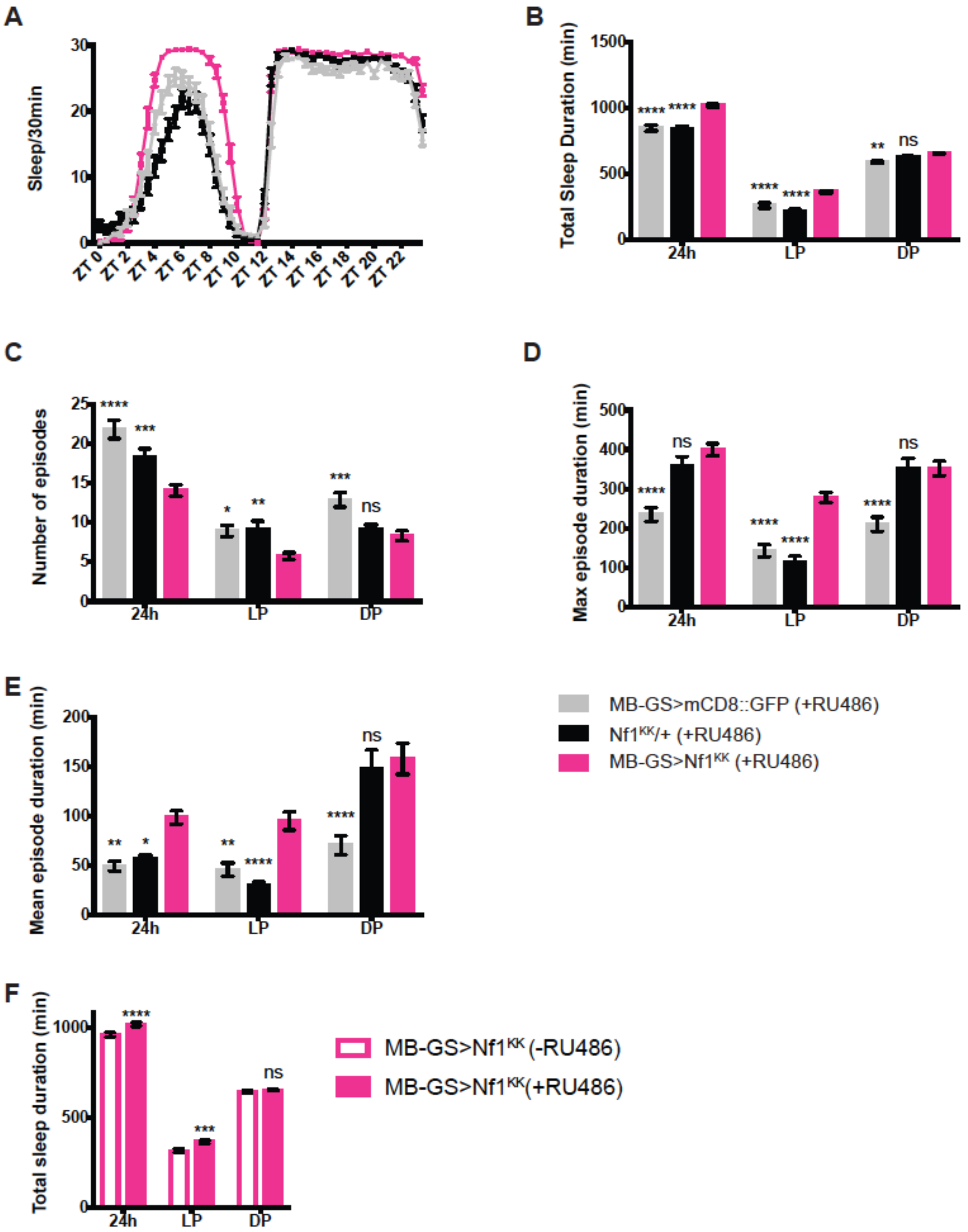


**Figure S2. NF1 in the adult MB promotes daytime wakefulness**

**A.** Sleep profiles of *MB-GS > NF1*$^{KK}$, *MB-GS > mCD8:GFP*, and *Nf1*$^{KK}$/+ fed with RU486 after eclosion to selectively induce MB-GS activity during adulthood. Sleep per 30-min bin averaged over 5 days of LD in the presence of RU486 is shown (n = 29–32 per group). **B−E.** Different sleep variables analyzed over 24 h, during the light phase (LP) and the dark phase (DP). All values represent the mean ± SEM. Values of *MB-GS > NF1*$^{KK}$ were compared with those of the two controls by two-way ANOVA with Dunnett's' correction for multiple comparisons ($*p < 0.05$, $**p < 0.01$, $***p < 0.001$, and $****p < 0.0001$). **B.** Total sleep duration. **C.** Number of sleep episodes. **D.** Maximum sleep episode duration. **E.** Mean sleep episode duration. **F.** Total sleep duration of the *MB-GS > NF1*$^{KK}$ flies fed with RU486 (+RU486) or those without RU486 (-RU486). Mean value ± SEM. ANOVA with Sidak's multiple comparison test found an increase in daytime sleep and sleep over a 24 h period following the induction of MB-GS with RU486 ($***p < 0.001$ and $****p < 0.0001$).

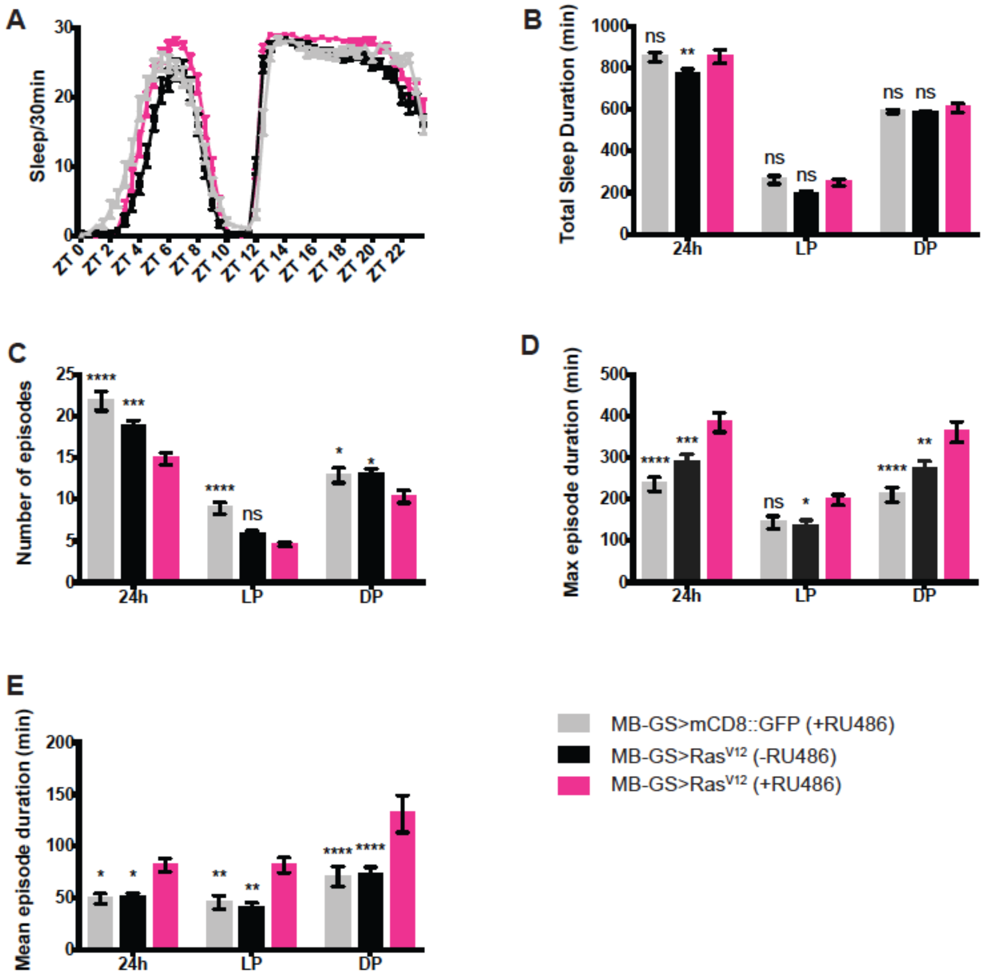


**Figure S3. Ras hyperactivation in the adult MB**

Sleep profiles, amount, and structure of *MB-GS* > *Ras*$^{v12}$ and *MB-GS* > *mCD8:GFP* flies fed with RU486 (+RU486) or those without RU486 (-RU486) during adulthood (n = 30 per group). All values represent the mean ± SEM. *MB-GS* > *Ras*$^{v12}$(+RU486) was compared with other groups by two-way ANOVA with Dunnett's multiple comparisons test. $*p < 0.05$, $**p < 0.01$, $***p < 0.001$, and $****p < 0.0001$). **A.** Sleep per 30-min bin averaged over 5 days of LD. **B.** Total sleep duration. **C.** Number of sleep episodes. **D.** Maximum sleep episode duration. **E.** Mean sleep episode duration.

**Table S1. Gene Ontology analysis of the MB LD-DD cyclers**
A total of 117 MB LD-DD cyclers were subjected to the GO term enrichment analysis using the GOrilla tool.

| GO term | Description | P-value | FDR q-value | Genes | |
|---|---|---|---|---|---|
| GO:0050890 | cognition | 2.37E-04 | 1.00E+00 | *CG1910* | *prt* |
| | | | | *w* | *Pka-C1* |
| | | | | *Nf1* | *CG4612* |
| | | | | *sgg* | |
| GO:0007611 | learning or memory | 2.37E-04 | 8.92E-01 | *CG1910* | *prt* |
| | | | | *w* | *Pka-C1* |
| | | | | *Nf1* | *CG4612* |
| | | | | *sgg* | |
| GO:0016070 | RNA metabolic process | 2.60E-04 | 6.51E-01 | *TfIIFbeta* | *Ars2* |
| | | | | *caz* | *CG4935* |
| | | | | *YT521-B* | *CG5800* |
| | | | | *CG2263* | *Mat1* |
| | | | | *CG16912 (α-PheRS)* | *CG11417* |
| | | | | *CG2972* | *CG4973*(*mdlc*) |
| | | | | *CG14450* | *wuho* |
| | | | | *Taf8* | *Clp* |
| | | | | *Ref1* | |
| GO:0008355 | olfactory learning | 2.89E-04 | 5.44E-01 | *prt* | *Pka-C1* |
| | | | | *Nf1* | *sgg* |
| GO:0008306 | associative learning | 6.59E-04 | 9.92E-01 | *prt* | *Pka-C1* |
| | | | | *Nf1* | *sgg* |
| GO:0015842 | aminergic neurotransmitter loading into synaptic vesicle | 7.02E-04 | 8.80E-01 | *prt* | *w* |
| GO:0015837 | amine transport | 7.02E-04 | 7.55E-01 | *prt* | *w* |
| GO:0006367 | transcription initiation from RNA polymerase II promoter | 8.35E-04 | 7.85E-01 | *TfIIFbeta* | *caz* |
| | | | | *Mat1* | *Taf8* |

**Table S2. List of the MB LD-DD cyclers selected for RNAi sleep screening**
LD adj.p and DD adj.p indicate adjusted *p*-values reported by JTK_Cycle in LD and DD, respectively. Gene snapshot describes the key information on the known function and biological roles of the gene.

| Flybase ID | Gene symbol | LD adj.p | DD adj.p | Gene snapshot |
|---|---|---|---|---|
| FBgn0000273 | *Pka−C1* | 1.34E-02 | 8.83E-03 | Protein kinase, cAMP-dependent, catalytic subunit 1. Axis specification, rhythmic behavior, synaptic transmission. |
| FBgn0003371 | *sgg* | 8.83E-03 | 4.30E-02 | Glycogen synthase kinase 3. A component of the β-catenin destruction complex in the canonical Wnt pathway. Circadian rhythms. |
| FBgn0015269 | *Nf1* | 4.30E-02 | 4.85E-04 | GTPase activating protein. Development, postembryonic growth, learning and memory, circadian rhythms. |
| FBgn0022349 | *CG1910* | 1.38E-03 | 1.34E-02 | Long-term memory. |
| FBgn0031414 | *eys* | 2.01E-02 | 8.83E-03 | Temperature entrainment of circadian rhythms. |
| FBgn0035016 | *CG4612* | 8.83E-03 | 5.71E-03 | Long-term memory, perception of pain. |
| FBgn0039714 | *Zip99C* | 3.62E-03 | 3.62E-03 | Zinc/iron regulated transporter-related protein 99C. |
| FBgn0043005 | *prt* | 3.62E-03 | 4.85E-04 | Vesicular transporter. Expressed in the mushroom body. |
| FBgn0259176 | *bun* | 8.83E-03 | 4.30E-02 | Mushroom body development, sleep. |
| FBgn0034029 | *eIF2Bγ* | 5.12E-07 | 3.62E-03 | Eukaryotic translation initiation factor 2B subunit gamma |
| FBgn0032484 | *kek4* | 4.30E-02 | 1.55E-04 | Transmembrane leucine-rich repeat (LRR) and immunoglobulin-like domain-containing (LIG) protein |
| FBgn0033309 | *CG8735* | 8.83E-03 | 8.83E-03 | Unknown |
| FBgn0037261 | *CG9775* | 3.62E-03 | 8.83E-03 | Unknown |
| FBgn0259984 | *kuz* | 2.01E-02 | 8.83E-03 | ADAM metalloendopeptidase. Axon guidance. |
| FBgn0037538 | *CG3223* | 2.01E-02 | 4.30E-02 | Unknown |
| FBgn0037051 | *CG10565* | 1.55E-04 | 1.55E-04 | Unknown |
| FBgn0029878 | *Pat1* | 2.01E-02 | 1.38E-03 | Regulation of microtubule-based movement. Localizes to the kinesin complex. |
| FBgn0040078 | *pont* | 2.01E-02 | 1.38E-03 | AAA+ (ATPases Associated with various cellular Activities) family. |
| FBgn0030007 | *α-PheRS* | 4.30E-02 | 1.55E-04 | Phenylalanyl-tRNA synthetase, α-subunit. |
| FBgn0035016 | *CG4612* | 8.83E-03 | 5.71E-03 | Predicted 3'UTR binding. Long-term memory. |
| FBgn0030871 | *CG8142* | 8.83E-03 | 8.83E-03 | Unknown |

**Table S3. Circadian locomotor rhythmicity of the flies following the knockdown of the MB LD-DD cyclers in the MB**

Some genotypes displayed reduced rhythmicity (*$p$ < 0.05, **$p$ < 0.01, ***$p$ < 0.001 according to the chi-square test, compared with *OK107-GAL4/+* ). No significant difference in the period (Tau ± SEM) was seen by knockdown as compared with *OK107-GAL4/+* flies (one-way ANOVA with Dunnett's multiple comparison test).

| Genotype | | % Rhythmic | Tau ± SEM (h) | Power ± SEM |
|---|---|---|---|---|
| *OK107>UAS-RNAi* | *Pka-C1* | 96.8 | 23.8 ± 0.05 | 161.2 ± 8.45 |
| | *Nf1* | 87.5 | 23.8 ± 0.04 | 153.8 ± 7.28 |
| | *CG1910* | 71.9* | 23.8 ± 0.05 | 116.1 ± 8.13 |
| | *eys* | 93.8 | 24.0 ± 0.02 | 161.6 ± 9.13 |
| | *zip99c* | 87.5 | 23.8 ± 0.05 | 149 ± 11.4 |
| | *portabella* | 100 | 23.9 ± 0.05 | 133.8 ± 8.11 |
| | *bun* | 90.6 | 24.2 ± 0.41 | 105.7 ± 7.13 |
| | *CG4616* | 81.5 | 24.3 ± 0.54 | 127.8 ± 10.18 |
| | *eIF2betgam* | 90.6 | 23.9 ± 0.03 | 165.9 ± 9.52 |
| | *hk2* | 87.5 | 23.9 ± 0.08 | 167.9 ± 18.25 |
| | *CG9775* | 73.7* | 23.2 ± 0.21 | 42.8 ± 4.62 |
| | *kuz* | 73.9* | 23.4 ± 0.09 | 36 ± 1.77 |
| | *CG3202* | 32.1*** | 23.4 ± 0.11 | 36.7 ± 5.98 |
| | *CG10565* | 57.7** | 23.2 ± 0.11 | 38.3 ± 3.45 |
| | *Dat1* | 70.6* | 23.4 ± 0.06 | 49 ± 4.19 |
| | *pont* | 36.4*** | 23.4 ± 0.13 | 40.6 ± 3.78 |
| | *alpha-PheRs* | 93.3 | 23.4 ± 0.12 | 42.4 ± 5.22 |
| | *CG8142* | 50*** | 23.3 ± 0.1 | 34.9 ± 2.77 |
| | *cabeza* | 44.8*** | 23.6 ± 0.1 | 33.6 ± 2.82 |
| | *kek-4* | 61.9*** | 23.5 ± 0.17 | 36.9 ± 3.54 |
| Controls | $w^{1118}$ | 96.9 | 23.4 ± 0.05 | 51.2 ± 3.43 |
| | *OK107/+* | 92.6 | 23.6 ± 0.05 | 662.3 ± 56.93 |

**Table S4. Effects on sleep induced by *Nf1* knockdown with different MB drivers**
Summary of the results displayed in Figure 6C. MB drivers expressed in different MB lobes were used to drive an RNAi transgene against *Nf1* (*UAS-*$Nf1^{KK}$). The column "Lobes" indicates the MB lobes where the corresponding driver is expressed. Total sleep duration in *GAL4 >* $Nf1^{KK}$ was compared with their driver-only and RNAi-only controls. The results are shown as compared with the GAL4-only control/ RNAi-only control. Sleep gain in *GAL4 >* $Nf1^{KK}$ relative to the control is marked as "Up" and sleep loss is marked as "Down". The significance level columns report the statistical test results using two-way ANOVA with Dunnett's correction for multiple comparisons (*$p < 0.05$, **$p < 0.01$, ***$p < 0.001$, and ****$p < 0.0001$). The first row (*OK107 >* $Nf1^{KK}$) is a summary of the data shown in Figure 3B. The comparisons with both controls showing significant differences in sleep gain or loss in the same direction are indicated in bold.

| Driver | lobes | 24h | | Light Phase | | Dark Phase | |
|---|---|---|---|---|---|---|---|
| | | Sleep change | Significance level | Sleep change | Significance level | Sleep change | Significance level |
| OK107 | all lobes | **Up/Up** | ****/**** | **Up/Up** | ****/**** | Up/Up | **/ns |
| c309 | γ, α/β, α′/β′ (weak) | Up/Up | ****/ns | Up/Up | ****/ns | Up/Down | ****/** |
| c747 | α/β, α′/β′, γ | Up/Down | ns/** | **Up/Up** | ****/**** | **Down/Down** | */**** |
| 30Y | α/β, α′/β′, γ | ****/Down | ns/ns | **Up/Up** | ****/* | Down/Down | ns/* |
| 201Y | γ and α/β (mostly core) | **Up/Up** | ****/**** | **Up/Up** | ****/**** | Up/Down | */* |
| 1471 | γ main | Up/Up | ns/ns | **Up/Up** | */**** | Down/Down | ns/** |
| mb247 | α/β, γ | Up/Up | ns/ns | **Up/Up** | ***/**** | Down/Down | ns/**** |
| 17D | α/β core | Up/Down | **/ns | Up/Up | ns/ns | Up/Down | */ns |
| c739 | α/β | Up/Up | ****/ns | **Up/Up** | */**** | Up/Down | */** |
| H24 | γ | Up/Down | ****/ns | Up/Up | **/ns | Up/Down | ****/ns |

**Table S5. Effects on sleep induced by *Nf1* knockdown with different KC split-GAL4 drivers**

Split-GAL4 drivers expressed in different KC subtypes were used to drive *UAS-Nf1*$^{KK}$. Total sleep duration (Total), number of sleep episodes (Episodes), maximum sleep episode duration (Max) and mean sleep episode duration (Mean) of *Split-GAL4 > Nf1*$^{KK}$ flies were compared with their driver-only and RNAi-only controls. The results are represented as described in Table S4. The significance columns report the test results using two-way ANOVA with Dunnett's correction for multiple comparisons ($*p < 0.05$, $**p < 0.01$, $***p < 0.001$, and $****p < 0.0001$; n = 26−55 per group).

| Driver | Lobes | Sleep parameter | 24h | | Light Phase | | Dark Phase | |
|---|---|---|---|---|---|---|---|---|
| | | | Sleep change | Significance | Sleep change | Significance | Sleep change | Significance |
| MB185B | α/βs | Total | Up/Up | ns/*** | Up/Up | ns/**** | Down/Down | ns/ns |
| | | Episode | Down/Up | */ns | **Down/Down** | ****/**** | Up/Up | ns/*** |
| | | Max | Up/Down | */ns | **Up/Up** | ******/****** | Up/Down | ns/**** |
| | | Mean | Up/Up | ns/ns | Up/Up | ns/** | Up/Down | ns/**** |
| MB594B | α/βc | Total | Down/UP | ns/** | Down/Up | ns/ns | Down/Up | ns/ns |
| | | Episode | **Up/Up** | ***/**** | Up/Up | ns/ns | Up/Up | ns/ns |
| | | Max | **Down/Down** | *****/*** | Down/Up | ns/ns | **Down/Down** | *****/**** |
| | | Mean | Down/Down | ns/ns | Down/Up | ns/ns | **Down/Down** | ****/**** |
| MB371B | α/βp | Total | Down/UP | ns/ns | Up/Up | ns/** | Down/Down | */ns |
| | | Episode | Up/Up | ns/*** | Down/Up | ns/ns | Up/Up | ns/**** |
| | | Max | Down/Down | ns/**** | Up/Up | ns/* | **Down/Down** | ***/****** |
| | | Mean | Down/Down | ns/ns | Up/Up | ns/ns | Down/Down | ns/**** |
| MB463B | α'/β'ap | Total | Down/UP | ns/**** | Up/Up | ns/*** | Down/Up | ns/ns |
| | | Episode | Up/Down | ns/ns | Down/Down | ns/ns | Up/Up | ns/ns |
| | | Max | Down/Down | **/ns | Up/Up | ns/** | Down/Down | ***/ns |
| | | Mean | Down/Up | ns/ns | Up/Up | ns/ns | Down/Down | ****/ns |
| MB418B | α'/β'm | Total | Up/Down | ns/ns | Up/Up | ns/ns | Up/Down | ns/* |
| | | Episode | Down/Up | */** | Down/Down | ns/ns | Down/Up | ns/* |
| | | Max | Down/Down | ns/**** | Down/Down | ns/ns | Down/Down | ns/**** |
| | | Mean | Up/Down | ns/ns | Up/Up | ns/ns | Up/Down | ns/**** |
| MB131B | γmain | Total | Up/Up | **/ns | Up/Up | ns/** | Up/Down | ns/ns |
| | | Episode | Down/Up | **/ns | Down/Down | **/ns | Up/Up | ns/* |
| | | Max | Up/Down | **/ns | **Up/Up** | *****/****** | Up/Down | ns/*** |
| | | Mean | Up/Down | ns/ns | Up/Up | ns/ns | Up/Down | ns/**** |
| MB607B | γd | Total | Up/Up | ns/*** | Up/Up | ns/*** | Down/Down | ns/ns |
| | | Episode | Down/Up | ns/* | Down/Down | ns/ns | Up/Up | ns/* |
| | | Max | Down/Down | ns/ns | **Up/Up** | ***/***** | Down/Down | ns/** |
| | | Mean | Up/Down | ns/ns | Up/Up | ns/ns | Down/Down | ns/*** |
